# Wnt-Activated Immunoregulatory Myeloid Cells Prevent Relapse in Experimental Autoimmune Encephalomyelitis and Offer a Potential Therapeutic Strategy for Multiple Sclerosis

**DOI:** 10.1101/2025.02.16.638560

**Authors:** Anastasia Alkhimovitch, Stephen D. Miller, Igal Ifergan

**Affiliations:** Division of Immunobiology, Cincinnati Children’s Hospital Medical Center, University of Cincinnati College of Medicine, Cincinnati, OH 45267; Department of Molecular and Cellular Biosciences, University of Cincinnati College of Medicine, Cincinnati, OH 45267; Neuroscience Graduate Program, University of Cincinnati College of Medicine, Cincinnati, OH 45229; Departments of Microbiology-Immunology, Feinberg School of Medicine, Northwestern University, Chicago, IL 60611; Departments of Interdepartmental Immunobiology Center, Feinberg School of Medicine, Northwestern University, Chicago, IL 60611

**Keywords:** Wnt signaling, Myeloid cells, Experimental autoimmune encephalomyelitis (EAE), Multiple sclerosis (MS), PD-L1, Immunoregulation

## Abstract

Multiple sclerosis (MS) is a chronic autoimmune disease of the central nervous system (CNS) characterized by recurrent inflammatory relapses and neurodegeneration. Myeloid cells play a critical role in shaping the inflammatory environment and influencing disease progression. Here, we demonstrate that activation of the Wnt signaling pathway reprograms myeloid cells into an immunoregulatory phenotype, leading to reduced neuroinflammation and disease severity. Using both experimental autoimmune encephalomyelitis (EAE) and human-derived myeloid cells, we show that Wnt agonist treatment promotes the expression of inhibitory molecules such as PD-L1 and PD-L2, suppressing pro-inflammatory responses. In the chronic and relapsing-remitting EAE models, Wnt activation significantly reduced disease severity, immune cell infiltration into the CNS, and pathogenic T cell responses. Notably, in relapsing-remitting EAE, Wnt treatment prevented new relapses in a PD-L1–dependent manner, highlighting the crucial role of myeloid cell–mediated immune regulation. These findings reveal a previously unrecognized role for Wnt signaling in myeloid cell immunoregulation and suggest that targeting this pathway could provide a novel therapeutic strategy for MS and other autoimmune diseases.

## Introduction

Multiple Sclerosis (MS) is a chronic autoimmune disease affecting approximately 2.8 million people worldwide (1). It is characterized by infiltration of peripheral immune cells into the central nervous system (CNS), leading to demyelination, neurodegeneration and progressive disability. Myeloid cells, including monocytes, dendritic cells (DCs) and macrophages are particularly abundant in CNS lesions in both MS and its widely used animal model, experimental autoimmune encephalomyelitis (EAE) (2–4). These cells function as antigen-presenting cells (APCs), facilitating T cell priming and expansion in secondary lymphoid organs and further sustaining T cell activation and cytokine polarization at sites of inflammation (5–7).

In MS/EAE, APCs exhibit increased antigen presentation capacity, primarily responsible for CD4^+^ T helper (Th)1 and Th17 activation and differentiation (8–11). Pro-inflammatory myeloid cells release inducible nitric oxide synthase (iNOS), reactive oxygen species, cytokines, and chemokines that exacerbate neurotoxicity in the CNS (12, 13). These inflammatory mediators further disrupt the blood-brain barrier (BBB) and promote the infiltration of additional peripheral immune systems into the CNS (14). During inflammation, APCs can secrete interleukin (IL)-12, a pivotal cytokine for CD4^+^ Th1 differentiation (15). Th1 cells, known for their production of interferon-gamma (IFN-γ), activate resident glial cells and recruit additional immune cells, thereby exacerbating CNS inflammation (16, 17). Moreover, APCs can produce IL-1β, IL-23, and IL-6, all of which are essential for Th17 differentiation (18). Th17 cells promote inflammation via the secretion of IL-17 and granulocyte-macrophage colony-stimulating factor (GM-CSF) (19, 20). IL-17 disrupts the BBB (16, 21), facilitating further immune infiltration, while GM-CSF activates myeloid cells and promotes an inflammatory profile in monocytes (22), establishing a feedback loop between the adaptive and innate immunity that drives chronic inflammation in MS (23).

In contrast, anti-inflammatory myeloid cells support tissue repair by promoting neuronal axon growth, clearing myelin debris, and facilitating remyelination (5, 6). These cells secrete anti-inflammatory cytokines such as IL-10, IL-4, and transforming growth factor-beta (TGFβ) (24, 25). The cytokines produced by these anti-inflammatory myeloid cells promote the differentiation of regulatory T cells (Tregs) (26, 27), which play a crucial role in suppressing Th1/Th17-driven inflammation, limiting disease severity in MS/EAE (28). While both pro- and anti-inflammatory myeloid cells play key roles in MS/EAE pathogenesis and recovery, current MS therapies primarily target lymphocytes, leaving the potential for myeloid-directed interventions largely unexplored.

The canonical Wnt/β-catenin (Wnt/β-cat) signaling pathway is critical for CNS homeostasis and immune regulation (29, 30). Wnt proteins are a family of lipoglycoproteins that bind to the cell surface Frizzled (Fzd) receptors, inhibiting glycogen synthase kinase-3β (GSK-3β) and preventing β-catenin (β-cat) degradation (31). This allows β-cat to accumulate and translocate to the nucleus where it activates the transcription of Wnt target genes via T cell factor (TCF) and lymphoid enhancer factor (LEF). Wnt/β-cat signaling has been implicated in immune modulation, particularly in inflammatory bowel disease (32), and colitis (33). Additionally, Wnt/β-cat activation in DCs promotes Tregs differentiation (34), while GSK-3β inhibition, an agonist of the Wnt/β-cat pathway, has been associated with neuroprotection in ischemic stroke (35) and traumatic brain injury (35, 36).

Despite these findings, the role of the Wnt/β-cat signaling in MS remains unclear. Some studies suggest that Wnt/β-cat activation in DCs suppresses neuroinflammation by modulating T cell responses, as reported by the Manicassamy group in chronic-(C-) EAE induced by MOG_35-55_

(37). In contrast, Yuan *et al.* proposed that upregulation of Wnt/β-cat signaling contributes to chronic pain in C-EAE (38). Given these discrepancies and the lack of research on Wnt signaling on myeloid cells during MS, we sought to determine whether this pathway modulates neuroinflammation in EAE. Furthermore, while previous studies have all focused on chronic MS models, our study explores the role of the Wnt/β-cat signaling in the relapsing-remitting form of MS (RRMS), which affects 80-85% of the patients (39). Understanding this pathway’s function in myeloid cells could provide new insights into MS pathogenesis and open avenues for novel therapeutic strategies.

## Material and Methods

### Mice and cell isolation

Female C57BL/6 mice and 2D2 breeder mice were purchased from The Jackson Laboratory (Bar Harbor, ME). Female SJL/J mice were obtained from Harlan Laboratories (Bethesda, MD). All mice were housed under specific pathogen-free conditions in the Association for Assessment and Accreditation of Laboratory Animal Care International (AAALAC)–approved facilities at Northwestern University and at the University of Cincinnati. CD11b^+^ myeloid cells, CD4^+^ T lymphocytes and CD4^+^ CD62L^hi^ CD44^lo^ naïve T lymphocytes were purified from C57BL/6 (for myeloid cells) or 2D2 (for T lymphocytes) spleens by magnetic cell sorting (Miltenyi Biotec, Auburn, CA) according to the manufacturer’s instructions.

### Human PBMCs and cell isolation

Peripheral blood mononuclear cell (PBMCs) suspensions of healthy donors (LifeSource, Evanston, IL) were obtained by density gradient centrifugation on Ficoll-Paque Plus (Cytiva Life Sciences). Human CD14^+^ monocytes, CD4^+^ CD45RA^+^ naïve T lymphocytes, and CD4^+^ CD45RA^−^ memory T lymphocytes were purified by magnetic cell sorting (Miltenyi Biotec) according to the manufacturer’s instructions.

### Activation of the Wnt/β-catenin pathway in vitro

To evaluate the effect of the Wnt agonist on human CD14^+^ monocytes, cells were plated at a density of 1.5 × 10^6^ cells/ml in 2ml in a 6 well plate in R10 media (RPMI with 10% (volume/volume) fetal bovine serum (FBS), 2 mM L-glutamine, 100 U/ml penicillin, 100 μg/ml streptomycin). For murine myeloid cells, CD11b^+^ cells were plated at a density of 1.5 – 2.5 × 10^6^ cells/ml in 200μl in a 96 well plate in R10 media. Cells were pre-treated with the Wnt agonist (SB216763) at a concentration of 10μM or with the vehicle control (dimethylsulfoxide (DMSO)) for 18 hours. For phosphorylated STAT and β-catenin analysis as well as quantitative PCR, a second stimulation with the Wnt agonist (10μM) or control vehicle was performed for 20min. After washing the media, cells were collected for either analysis or plated again for co-culture with CD4^+^ T lymphocytes.

### Human mixed leukocyte reaction and 2D2 lymphocyte activation

Wnt-activated or vehicle control cells were washed twice with PBS, collected, counted and plated in 96-well plates. For human mixed leukocyte reaction, allogeneic naïve and memory CD4^+^ T lymphocytes (5 × 10^5^ cells/well) were added to monocytes (1 × 10^5^ cells/well) and cells were incubated for 5 days in R10 media. For 2D2 lymphocyte activation, 5 × 10^5^ naïve or total CD4^+^ T lymphocytes were plated in presence of 0.5 × 10^5^ CD11b^+^ cells with MOG_35-55_ for 4 days in R10 media. CD4 profile, cytokine expression and proliferation were assessed by flow cytometry. Proliferation was evaluated after staining for Ki-67 (Thermo Fisher Scientific) nuclear antigen. Live-dead discrimination was performed using LIVE/DEAD fixable staining reagents (Life Technologies) and intracellular staining for Ki-67 was done using Foxp3/Transcription Factor Staining Buffer Set (Thermo Fisher Scientific). The frequency of Ki-67 positive cells was assessed on gated live CD3^+^ CD4^+^ cells. For cytokine secretion, media samples were measured by multiplex cytokine assays (Millipore) for IFN-γ, IL-17, GM-CSF, TNF and IL-10 production according to manufacturer’s instructions.

### Gene expression profile

Wnt-activated or vehicle control monocytes (n = 5 donors) were washed twice with PBS and lysed using 350μl of buffer RLT (Qiagen). RNA was isolated using a RNeasy Mini kit (Qiagen) following the manufacturer’s instructions. Purity of the isolated RNA was determined by measuring the ratio of the optical density of the samples at 260 and 280 nm using a Nanodrop spectrophotometer. The OD260/OD280 ratio ranged from 1.9 to 2.1 for all samples. cDNAs were synthesized using RT2 First Strand kit (Qiagen), according to the manufacturer’s instructions. The Human JAK/STAT Signaling Pathway RT2 Profiler PCR Arrays were purchased from SABiosciences, Qiagen. This array profiles the expression of 84 genes of the Jak/STAT pathway and includes 5 controls for housekeeping genes, one control for genomic DNA and three reverse transcription controls. PCRs were performed on a QuantStudio 3. The data were analyzed using the web-based software RT2 Profiler PCR Array data analysis tool (Qiagen). Fold-changes for each gene were calculated as the difference in gene expression between the vehicle-treated monocytes and Wnt agonist-treated monocytes. A positive value indicates a gene up-regulated in the Wnt agonist-treated monocytes and a negative value indicates a gene down-regulated in the Wnt agonist-treated monocytes.

### β-catenin and phosphorylated STAT ELISA

Analysis of total β-catenin, phosphorylated β-catenin, and phosphorylated STAT were assessed from cell lysates using InstantOne enzyme-linked immunosorbent assay (ELISA) kits (Thermo Fisher Scientific), according to the manufacturer’s protocol. Plates were read at 450 nm wavelength and analyzed using SoftMax Pro 5.2 program (Molecular Devices). Results are expressed as the mean absorbance of duplicate wells.

### Active EAE

Eight- to ten-week-old female C57BL/6 mice were used to induce EAE by active immunization. Mice were injected subcutaneously with 200 μg of MOG_35-55_ peptide (Genemed Synthesis, San Francisco, CA) emulsified in complete Freund’s adjuvant (CFA) containing 200 μg of *Mycobacterium tuberculosis* H37Ra (Difco, Detroit, MI) distributed over three sites on the flank. On day 0 and 2 after immunization, 200 ng pertussis toxin (List Biological Laboratories, Campbell, CA) were administered intraperitoneally (i.p.). Mice were treated with the Wnt agonist (SB216763 from Tocris Bioscience; 0.5 mg/kg, 2.5 mg/kg or 10 mg/kg) resuspended in dimethylsulfoxide (DMSO) and further diluted with the Tocrisolve 100 (from Tocris Bioscience), or control vehicle which consisted of DMSO diluted with the Tocrisolve 100. Injections started at day -7 and were done i.p. three times a week, until day 11, when control mice showed signs of disease. Following the initial experiment testing doses, the ensuing experiments were performed using 2.5 mg/kg. EAE experiments using C57BL/6 mice were performed 3 times with 8-12 mice for each of them.

For RR-EAE disease induction, eight- to ten-week-old female SJL/J mice were injected with CFA emulsion containing 50 μg PLP_139–151_. Mice were treated the Wnt agonist (2.5 mg/kg) or the vehicle control after the first relapse (day 34 post induction). The injections were done i.p. three times a week, until the completion of the study. Two independent experiments were performed, both experiments with 8-12 mice per group.

Treatments with anti-PD-L1 (clone 10F.9G2 from Bioxcell; 10mg/kg) or anti-IL-10 (clone JES5-2A5 from Bioxcell; 10mg/kg) antibodies or isotype control were injected at the same time as the Wnt agonist treatment started, and were done i.p. three times a week, for two consecutive weeks.

Clinical signs of EAE were assessed daily according to the following scores: 0, no clinical sign of disease; 1, limp tail; 2, hind limp weakness; 3, partial hind limb paralysis; 4, complete hind limb paralysis; 5, hind and fore limb paralysis. Data are reported as the mean daily clinical score.

### Ex vivo recall responses and LPS activation of splenocytes

Spleens were harvested from mice at day 14 post-immunization (for EAE induced on C57BL/6 mice) or day 62 post-immunization (for EAE induced on SJL/J mice), counted, and cultured in 96-well microtiter plates at a density of 10^6^ cells/well in a total volume of 200 μl of R10 media. Cells were cultured at 37°C in the presence of OVA_323–339_ (purity: 97.29%), PLP_139–151_ (purity: 97.78%), PLP_178–191_ (purity: 95.12%), MOG_35-55_ (purity: 98.41%), MBP_84-97_ (purity: 96.34%) (Genemed Synthesis; 20 μg/ml) or in the absence of peptide. Forty-eight hours post-culture initiation, the wells were pulsed with 1 μCi of [^3^H]-TdR, and the cultures were harvested on day 3. Results are expressed as the mean cpm of triplicate cultures. For cytokine quantification, media samples were measured by multiplex cytokine assays (Millipore) for IFN-γ, IL-17, GM-CSF and TNF-α production according to manufacturer’s instructions. In addition, CD4^+^ T cells profile was analyzed by flow cytometry. Expression of PD-1, CTLA-4, CD25, FoxP3 and TGF-β1 was assessed following the recall responses.

Splenocytes (10^6^ cells/well) were also activated for 24h in presence of LPS (10 ng/ml) from *E. coli* serotype 0111:B4 (Sigma) in 200 μl of R10 media. IL-1β, IL-6, IL-10, IL-12p70, IL-23 and TNF-α cytokines were measured by multiplex cytokine assays (Millipore) following manufacturer’s instructions.

### Isolation of CNS leukocytes

CNS-immune cells were isolated by Percoll gradient centrifugation from homogenized combined brain and spinal cords as previously described (40). The numbers of cell subpopulations in the CNS were determined by multiplying the percentage of lineage marker– positive cells by the total number of mononuclear cells isolated from the CNS.

### Flow Cytometry Analysis

In order to carry out flow cytometry analysis, the Fc receptors were initially blocked using anti-mouse CD16/32 (0.25 μg; Thermo Fisher Scientific) for 15 min at 4°C. Cells were then washed with fluorescence-activated cell sorting (FACS) buffer containing PBS with 1% (v/v) fetal bovine serum and 0.1% (w/v) NaN3 (Sigma). Cells were then stained for surface marker for 30 min at 4°C using the specified antibodies. To detect T cell cytokine expression, cells were activated for 18 h with 1 mg/ml ionomycin and 20 ng/ml phorbol 12-myristate 13-acetate 40 (PMA) in the presence of 2 mg/ml brefeldin A (Sigma) for the last 6 h of co-culture. To detect myeloid cell cytokine expression, cells were activated for 18 h with 100 ng/ml LPS from E. coli serotype 0111:B4 (Sigma) in the presence of 2 mg/ml brefeldin A (Sigma) for the last 6 h of co-culture. Cells were stained for surface markers: CD45, CD11b, CD11c, Ly6C, Ly6G, MHC II, CD40, CD80, CD86, PD-L1, PD-L2, CD3, CD4, CD25, PD-1 and TGF-β1 (all from Biolegend, BD Biosciences and Thermo Fisher Scientific). Cells were then washed with PBS and viability staining was then performed using the LIVE/DEAD fixable dead cell stain kit (Invitrogen). Following viability staining, cells were washed with PBS and were either resuspended in FACS buffer for Flow cytometry analysis or were used for intracellular staining to detect IL-1β, IL-6, IL-10, IL-12p40, IL-23p19, TNF, IL-17, IFN-γ, GM-CSF, CTLA-4 (CD152), FoxP3 and Ki-67. For intracellular staining, cells were fixed and permeabilized using the FoxP3 staining buffer kit (Thermo Fisher Scientific), and then intracellularly stained. Cells were acquired on a BD Canto II and analyzed using BD FACSDiva version 6.1 software. As controls, fluorescence minus one (FMOs) were used to place the gates for analysis.

For phosphorylated STAT staining, after stimulation with the Wnt agonist or the control vehicle for 18h, media was washed twice with PBS, and live-dead staining was performed using LIVE/DEAD fixable staining reagents (Life Technologies). Cells were stimulated a second time with the Wnt agonist or the control vehicle for 20min in R10 media. Cells were then fixed by adding BD Cytofix Fixation Buffer (BD Biosciences) and incubating at 37°C for 10min. After centrifugation, cells were permeabilized by adding BD Phosflow Perm Buffer III (BD Biosciences) and incubating for 30 minutes on ice. After 2 washes with FACS buffer, cells were then stained with pSTAT antibodies.

For myeloid cell analysis, cells were first gated according to FSC-SSC, then restricted to singles cells and live cells. In the CNS, infiltrating myeloid cells were identified as CD45^hi^ CD11b^+^. Ly6G^+^ neutrophils were first gated and excluded from the myeloid sub-populations. The Ly6G^−^ myeloid cells were divided into CD11c^+^ mDCs and CD11c^−^ monocytes/macrophages. Finally, the monocytes/macrophages were further divided into Ly6C^hi^ inflammatory monocytes, and Ly6c^lo^ non-inflammatory monocytes.

### Statistical analysis

Statistical analyses were performed using GraphPad PRISM 9.3 (GraphPad software). Data are presented as the mean ± the standard error of the mean (SEM). All the in vitro analyses comparing the effect of the Wnt agonist to the vehicle control were performed by paired T-test. When more than 2 variables were present, 2-way ANOVA tests were performed. EAE scores were analyzed by nonparametric Mann-Whitney test. Only p values < 0.05 were considered significant.

## Results

### Wnt/β-catenin activation drives an anti-inflammatory profile in human monocytes

To investigate the effect of Wnt/β-catenin activation on human APCs, we isolated CD14^+^ monocytes from healthy donors. These cells were treated with the Wnt agonist, SB216763, at a concentration of 10μM. SB216763 inhibits the enzyme GSK3β, thereby enabling constitutive activation of the canonical Wnt pathway.

Our first objective was to confirm that this concentration of the Wnt agonist effectively activated the Wnt/β-catenin pathway. To do so, we quantified total β-catenin and phosphorylated β-catenin levels in monocytes. We observed a significant increase in total β-catenin following treatment with the Wnt agonist (**Figure S1**; n = 6 donors, \*\*\**p* < 0.001), whereas phosphorylated β-catenin levels remained unchanged (**Figure S1**; n = 6 donors, not significant). These results indicate that our chosen concentration of SB216763 allows β-catenin accumulation without increasing its phosphorylated, thereby preventing its degradation. These findings confirm the successful activation of the canonical Wnt pathway.

Following the confirmation of our activation protocol, we next analyzed the immunological profile of human monocytes after Wnt/β-catenin activation. First, we examined the expression of costimulatory molecules, including CD80, CD86 and CD40. Monocytes showed no significant changes in the expression of these molecules following Wnt agonist treatment (**Figure 1A upper panels and Figure S2;** n = 7 donors). We then assessed the expression of inhibitory molecules PD-L1 and PD-L2 and found a significant upregulation after Wnt/β-catenin activation (**Figure 1A lower panels** and **Figure S2**; n = 7 donors, \*\**p* < 0.01, \*\*\**p* < 0.001).

**Figure 1.**
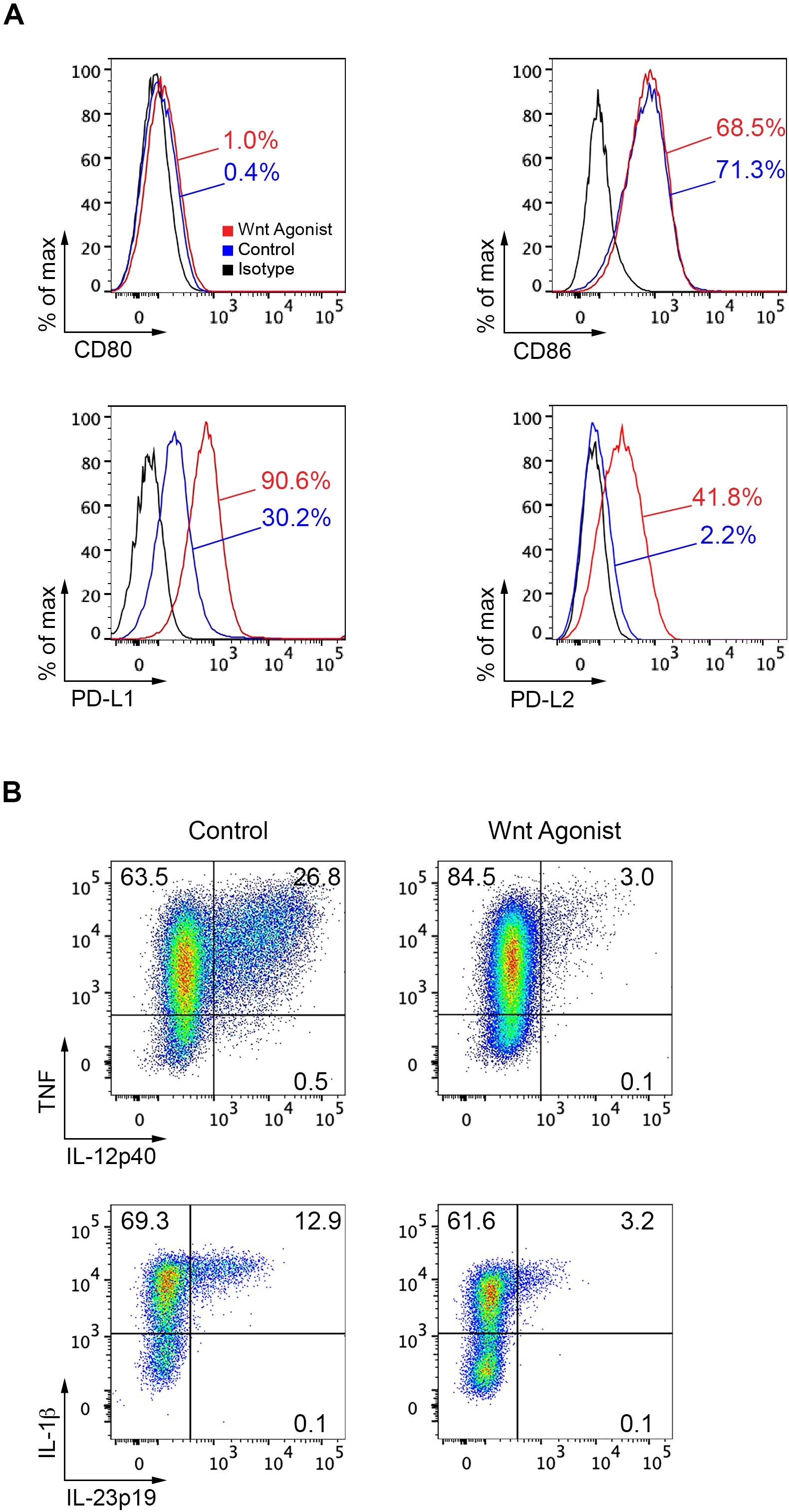
Activation of Wnt/β-Catenin Pathway Drives an Anti-Inflammatory Profile in Human Monocytes. CD14^+^ Monocytes were isolated from the blood of healthy donors and treated with either a Wnt agonist (SB216763, 10μM) or vehicle (DMSO) for 18 hours. Cells were then washed, and flow cytometry analysis was performed. (**A**) Representative flow cytometry plots showing expression of costimulatory (CD80, CD86) and inhibitory (PD-L1, PD-L2) molecules in human CD14⁺ monocytes. Cells treated with Wnt agonist are shown in red, vehicle-treated cells in blue, and the isotype staining in black. (**B**) Flow cytometry plots of IL-12p40, IL-23p19, and IL-1β expression in Wnt-treated (right) vs. control monocytes (left). Data shown are representative of 7 independent experiments.

Next, we evaluated cytokine expression in human monocytes and observed a significant downregulation of the pro-inflammatory cytokines IL-12p40, IL-23p19 and IL-1β in Wnt-activated monocytes compared to control monocytes (**Figure 1B and Figure S2**; n = 7 donors, \**p* < 0.05, \*\**p* < 0.01, \*\*\**p* < 0.001). No significant differences were found for other cytokines (**Figure S2**). Notably, in contrast to previous findings (PMID: 25710911), IL-10 expression was not upregulated upon Wnt activation (**Figure S2**).

To further characterize the immunoregulatory profile of Wnt-activated monocytes, we performed qPCR analysis of CD14^+^ monocytes culture in the presence or absence of the Wnt agonist. We observed a significant downregulation of Nos2 (also denoted iNOS) gene expression following Wnt/β-catenin activation (**Figure 2A and Table S1**; n = 6 donors, \**p* < 0.05). Since iNOS is a key marker of monocytes/macrophages activation, its reduced expression indicates a shift toward a less inflammatory phenotype. Collectively, these data demonstrate that Wnt pathway activation drives human monocytes toward a regulatory phenotype.

**Figure 2.**
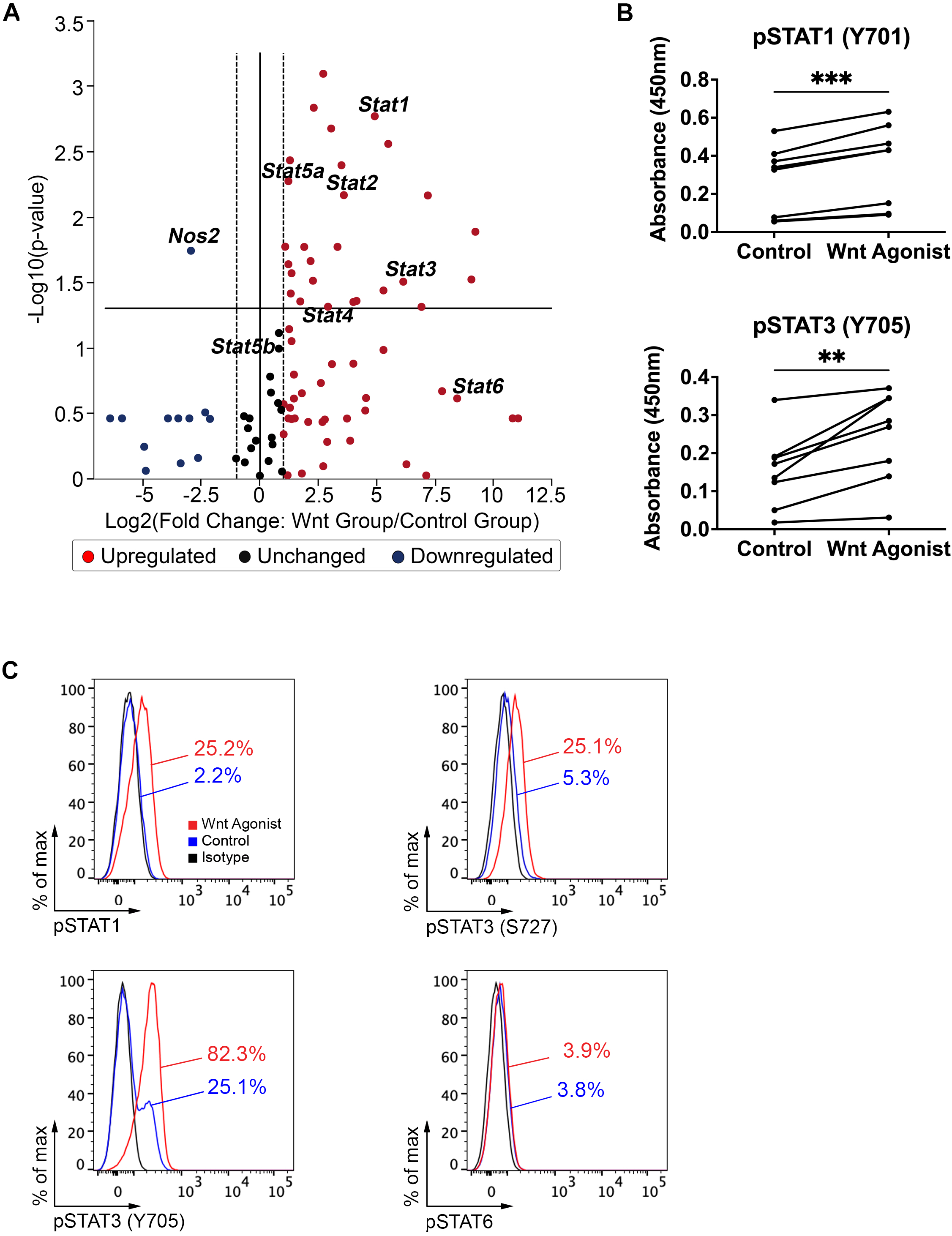
Wnt/β-Catenin Activation Induces STAT Signaling in Human Monocytes. CD14^+^ Monocytes were isolated from the blood of healthy donors and treated with either a Wnt agonist (SB216763, 10μM) or vehicle (DMSO) for 18 hours. A second stimulation with the Wnt agonist (10μM) or control vehicle (DMSO) was performed for 20min for qPCR and phosphorylated STAT (pSTAT) analysis. (**A**) Volcano plot of differential gene expression by qPCR, showing statistical significance vs. fold change. Each point represents a gene, where the x-axis indicates the log2 fold change for Wnt-treated monocytes vs. vehicle-treated monocytes, and the y-axis represents the –log10 of the *p*-value. Vertical dashed lines indicate ± 1.5-fold changes and the horizontal line marks a *p*-value threshold of 0.05 (Student’s *t*-test, *n* = 6 donors). A full list of genes and *p*-values is provided in the Supporting Material (**Table S1**). (**B**) ELISA quantification of pSTAT1 and pSTAT3 (Y705) following Wnt activation (*n* = 8 donors, \*\**p* < 0.01, \*\*\**p* < 0.001 by paired Student’s *t* test). (**C**) Representative histograms from flow cytometry analysis showing expression of pSTAT1, pSTAT3 (S727), pSTAT3 (Y705), pSTAT6 in Wnt-treated monocytes compared to vehicle control. Wnt-treated cells are in red, vehicle-treated cells in blue, and the isotype staining in black. Data shown are representative of 8 independent experiments.

### Wnt/β-catenin activation induces STAT signaling in human monocytes

There have been limited reports indicating that PD-L1 and PD-L2 can be upregulated following Wnt activation (PMID: 34781737; PMID: 30705400). However, the molecular mechanisms underlying this regulation remain unclear. Signal transducer and activator of transcription (STATs) family members have been implicated in Wnt signaling in various cell types (41–43) and are known to regulate PD-L1 and PD-L2 expression (44–46). Therefore, we examined STAT expression using qPCR, and found that *Stat1, 2, 3, 4 and 5a* were upregulated in Wnt-activated human monocytes (**Figure 2A and Table S1**; n = 6 donors, \**p* < 0.05).

To further explore STAT activation, we measured the expression of phosphorylated STAT (p-STAT) proteins by ELISA and flow cytometry. STAT proteins are phosphorylated by Janus kinase (JAK) family members, allowing their translocation to the nucleus, where they regulate gene transcription. Upon Wnt/β-catenin activation, we detected increased expression of p-STAT1, p-STAT3 (S727) and p-STAT3 (Y705), all of which have been linked to PD-L1 regulation (46) (**Figure 2B and 2C** and **Figure S3**; n = 8 donors, \**\*p* < 0.01, \*\*\**p* < 0.001). In contrast, PD-L2 expression is known to be regulated by pSTAT1 and pSTAT6 (45). However, only pSTAT1 was upregulated following Wnt activation. Taken together, these results suggest that Wnt/β-catenin activation promotes PD-L1 and PD-L2 expression through upregulation of the STAT signaling pathways.

### Inhibition of CD4 inflammatory response by Wnt-activated monocytes

We next assessed the ability of Wnt-activated monocytes to influence T cell activation. To this end, we performed a mixed leukocyte reaction (MLR) and measured CD4^+^ T cell proliferation and cytokines. Human CD4^+^ CD45RA^+^ naïve T cells and CD4^+^ CD45RA^−^ memory T cells were isolated and co-culture with either Wnt-activated or control monocytes. Naïve CD4^+^ T cells exhibited increased proliferation when co-culture with Wnt-activated monocytes compared to control monocytes (**Figure 3A right panel**; n = 11 MLRs, \*\**p* < 0.01). In contrast, memory CD4^+^ T cell proliferation remained unchanged between the two conditions (**Figure 3A, left panel**). However, cytokine analysis revealed that memory CD4^+^ T cells co-cultured with Wnt-activated monocytes expressed significantly lower levels of pro-inflammatory cytokines, including IL-17, IFN-γ, GM-CSF and TNF, compared to those co-cultured with control-monocytes (**Figure 3B**; n = 11 MLRs, \**p* < 0.05, \*\**p* < 0.01).

**Figure 3.**
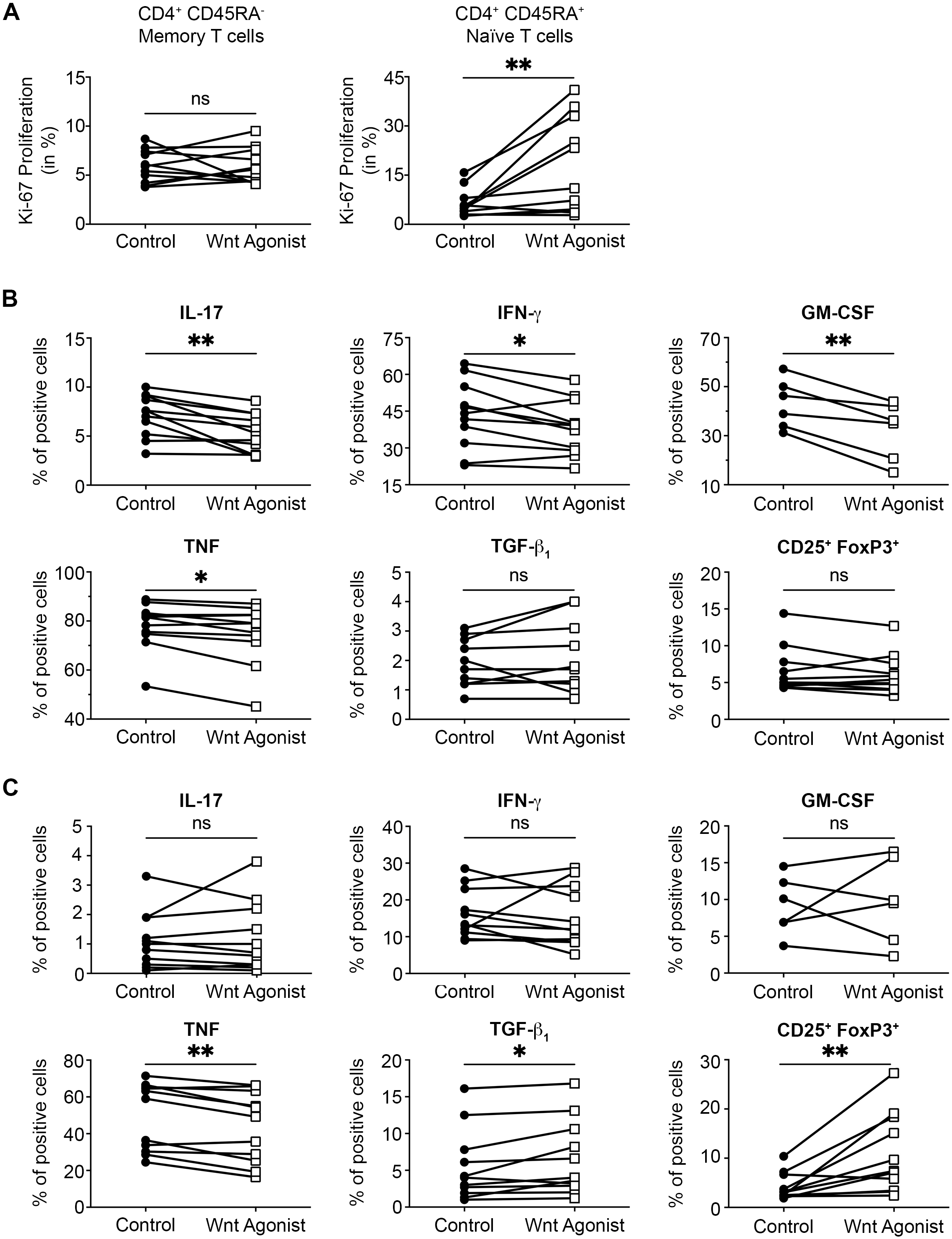
Wnt-Activated Monocytes Suppress Inflammatory CD4^+^ T Cell Responses. Human CD4^+^ T lymphocytes were co-cultured in a mixed leukocyte reaction (MLR) with Wnt-treated (SB216763, 10μM; open squares) or control (DMSO; closed circles) monocytes for 5 days. (**A**) Flow cytometry analysis of CD4⁺ T cell proliferation assessed by % of Ki-67^+^. Naïve CD4^+^ CD45RA^+^ T cells showed increased proliferation, whereas memory CD4^+^ CD45RA^−^ T cell proliferation remained unchanged (*n* = 11 MLRs, \*\**p* < 0.01 and ns = not significant by paired Student’s *t* test). (**B**) Cytokine quantification via flow cytometry from memory CD4^+^ CD45RA^−^ T cell co-cultured with Wnt-activated monocytes, showing decreased IL-17, IFN-γ, GM-CSF, and TNF expression (*n* = 11 MLRs, \**p* < 0.05, \*\**p* < 0.01 and ns = not significant by paired Student’s *t* test). (**C**) Cytokine quantification via flow cytometry from naïve CD4^+^ CD45RA^+^ T cells co-cultures with Wnt-activated monocytes, showing increased TNF and TGF-β, along with enhanced regulatory T cell (Treg) differentiation (CD25⁺ FoxP3⁺) (*n* = 11 MLRs, \**p* < 0.05, \*\**p* < 0.01 and ns = not significant by paired Student’s *t* test).

Analysis of the co-culture supernatants further supported these findings, showing significant reduction in the levels of inflammatory cytokines IFN-γ and GM-CSF, while the anti-inflammatory cytokine IL-10 in memory CD4^+^ T cells co-cultured with Wnt-activated monocytes were shown to be elevated (**Figure S4A**; n = 11 MLRs, \**p* < 0.05, \*\**p* < 0.01). Co-culture of Wnt-activated monocytes with naïve CD4^+^ T cells resulted in reduced TNF expression, increased TGF-β expression, elevated IL-10 secretion, and enhanced polarization towards CD25^+^ FoxP3^+^ regulatory T cells (Tregs) compared to co-culture with control-monocytes (**Figure 3C and Figure S4B**; n = 11 MLRs, \**p* < 0.05, \*\**p* < 0.01). These findings suggest that Wnt-activated monocytes promote a regulatory T cell response during the priming phase and decease inflammatory T cell responses during the reactivation phase.

### Murine CD11b^+^ myeloid cells recapitulate the Human Wnt-mediated regulatory profile

To validate these findings in a murine model, we investigated whether Wnt/β-catenin activation in mouse APCs induces a similar regulatory phenotype. We isolated CD11b^+^ myeloid cells from the spleen of healthy mice and exposed them to the Wnt agonist SB216763 for 24h. Consistent with our human data, Wnt-activated murine CD11b^+^ showed significantly increased PD-L1 and PD-L2 expression while downregulating their IL-12p40 and IL-23p19 expression (**Figure 4A and Figure S5**; n = 5, \**p* < 0.05, \*\**p* < 0.01). Analysis of STAT activation revealed increase in pSTAT1, pSTAT3 (S727) and pSTAT3 (Y705) following Wnt pathway activation in CD11b^+^ cells (**Figure 4B**; n = 5, \**p* < 0.05, \*\**p* < 0.01, \*\*\**p* < 0.001).

**Figure 4.**
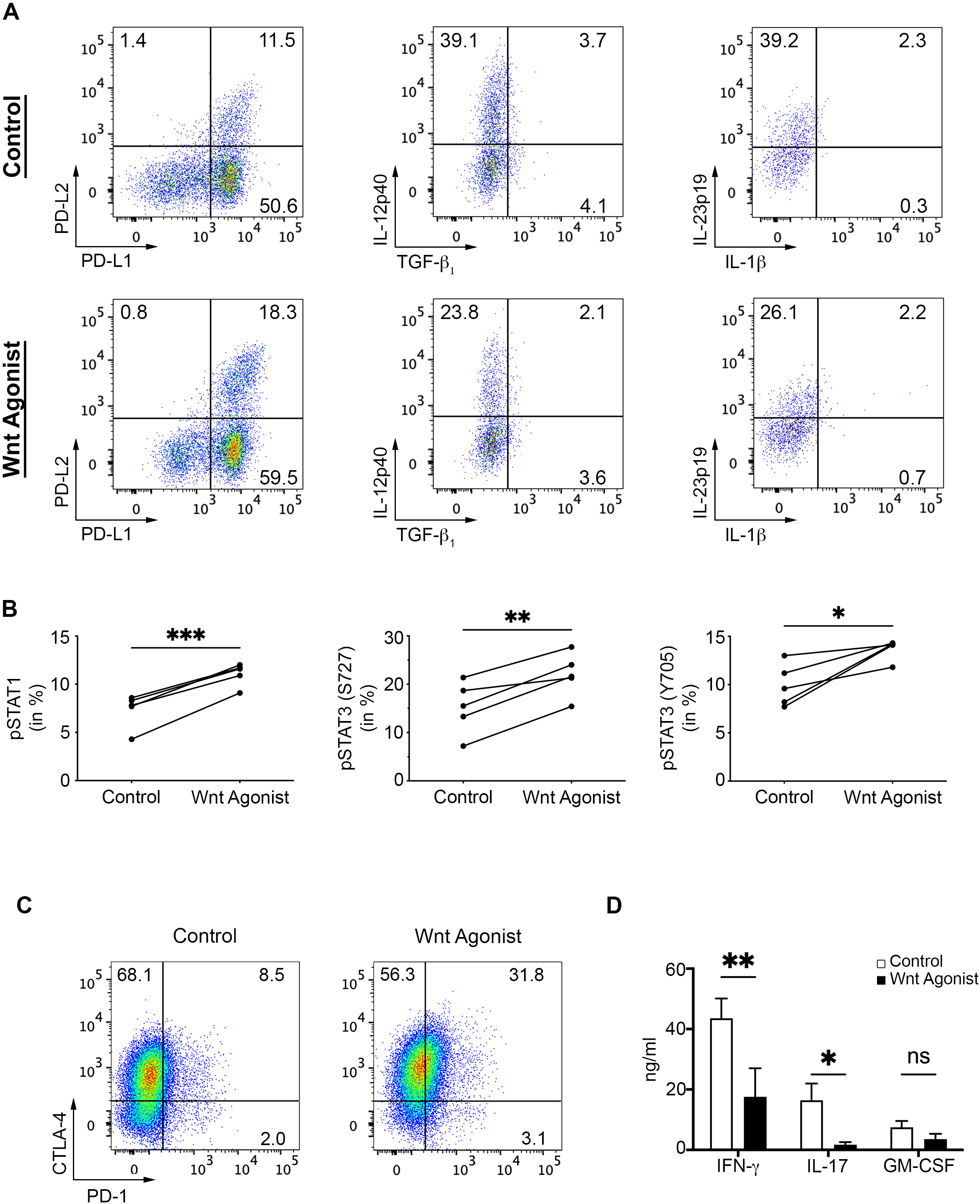
Murine Myeloid Cells Recapitulate the Human Wnt-Mediated Regulatory Profile. CD11b^+^ splenocytes were isolated from healthy adult C57BL/6 mice and treated with either a Wnt agonist (SB216763, 10μM) or vehicle (DMSO) for 24 hours. Cells were then washed, and flow cytometry analysis was performed. (**A**) Representative flow cytometry plots showing expression of inhibitory (PD-L1, PD-L2) molecules and cytokines (IL-12p40, TGF-β, IL-1β and IL-23p19) in murine CD11b⁺ myeloid cells. Data shown are representative of 5 independent experiments. (**B**) ELISA analysis showing increased pSTAT1, pSTAT3 (S727), and pSTAT3 (Y705) in Wnt-treated murine CD11b⁺ cells (*n* = 5, \**p* < 0.05, \*\**p* < 0.01, \*\*\**p* < 0.001 by paired Student’s *t* test). (**C**) Representative flow cytometry plots showing increased expression of CTLA-4 and PD-1 in naïve CD4^+^ CD62L^hi^ CD44^lo^ T cells isolated from the spleen of healthy adult 2D2 mice co-cultured with either Wnt-activated (right panel) or vehicle (left panel) murine CD11b⁺ cells for 4 days. Data shown are representative of 5 independent experiments. (**D**) Multiplex cytokine assay analysis showing decreased IFN-γ and IL-17 secretion by total 2D2 CD4⁺ T cells cultured with Wnt-treated CD11b⁺ cells (black bar) compared to vehicle control CD11b⁺ cells (white bar) (*n* = 5, \**p* < 0.05, \*\**p* < 0.01 and ns = not significant by two-way ANOVA with a Bonferroni post-test).

Next, we examined the antigen presentation capacity of Wnt-activated murine myeloid cells. CD62L^hi^ CD44^lo^ naïve CD4^+^ T cells and total CD4^+^ T cells were isolated from 2D2 transgenic mice, which express a TCR specific to MOG_35-55_. These T cells were co-cultured with Wnt-activated or control-CD11b^+^ cells in the presence of the MOG_35-55_ peptide. Similar to our human data, naïve CD4^+^ T cells showed significantly increased proliferation when co-cultured with Wnt-activated CD11b^+^ cells (**Figure S6, top panel**; n = 5, \*\**p* < 0.01). However, unlike in human MLR assays, murine naïve CD4^+^ T cells did not show an increase in Tregs differentiation following co-culture with Wnt-activated CD11b^+^ cells (**Figure S6)**. Instead, these cells significantly upregulated the expression of the immunoregulatory molecules CTLA-4 and PD-1 compared to those cultured with control CD11b^+^ cells (**Figure 4C and Figure S6**; n = 5, \*\**p* < 0.01). No significant differences in cytokine expression were observed in naïve CD4^+^ T cells (**Figure S6**).

Regarding total CD4^+^ T cells, we observed a significant decrease in IFN-γ and IL-17 secretion when co-cultured with Wnt-activated CD11b^+^ cells (**Figure 4D**, n = 5, \**p* < 0.05, \*\**p* < 0.01), whereas no changes in proliferation were detected (data not shown). Collectively, these results confirm our human findings and demonstrate that Wnt pathway activation in both human and murine myeloid cells induces an immunoregulatory phenotype, which in turn modulates CD4^+^ T cell inflammatory potential.

### Prophylactic treatment with a Wnt agonist ameliorates chronic EAE

Myeloid cells constitute a significant proportion of leukocytes found in MS and EAE lesions. These cells play a pivotal role in engulfing myelin debris and presenting antigen to T cells, thereby influencing their activation and differentiation (5–7). To evaluate the capacity of Wnt activation in modulating CNS inflammation, we induced C-EAE with MOG_35-55_ in C57BL/6 mice. The goal was to administer the Wnt agonist multiple times per week to ensure Wnt activation of newly generated circulating myeloid cells.

Given our *in vitro* findings, we hypothesized that Wnt-activated myeloid cells could influence T cell priming. Thus, in our initial *in vivo* experiment, Wnt agonist administration started seven days before disease induction (day -7) and lasted until day 11 post-immunization, when T cell priming was completed.

To our knowledge, repeated dosing of a Wnt agonist in animals has not been previously explored. Therefore, we tested three different doses, 0.5, 2.5 and 10mg/kg/mouse, 3 times a week, through the i.p. route. At doses of 2.5 and 10mg/kg, we observed a significant reduction in disease severity compared to vehicle controls, whereas the 0.5mg/kg had no significant effect (**Figure S7**; n = 5 mice per group, \**p* < 0.05). Since no significant difference was observed between 2.5mg/kg and 10mg/kg groups, we selected 2.5mg/kg for all subsequent experiments.

We repeated the experiment, again administrating the Wnt agonist at 2.5mg/kg 3 times a week from day-7 until day 11 via i.p. administration. We again observed a significant reduction in disease severity (**Figure 5A**; n = 10 mice per group, \*\*\**p* < 0.001). Analysis of CNS infiltrating immune cells at day 18 post-immunization revealed that Wnt agonist treatment significantly reduced the total number of CNS infiltrating cells, the number of CD45^hi^ CD3^−^ CD11b^+^ infiltrating myeloid cells, and of CD45^hi^ CD3^+^ CD4^+^ T lymphocytes (**Figure 5B**; n = 5 mice per group, *\*p* < 0.05). Furthermore, mice treated with the Wnt agonist exhibited a significant reduction in CD4^+^ T lymphocytes expressing IFN-γ, IL-17, GM-CSF and TNF (**Figure 5C**; n = 5 mice per group, *\*p* < 0.05, *\*\*p* < 0.01).

**Figure 5.**
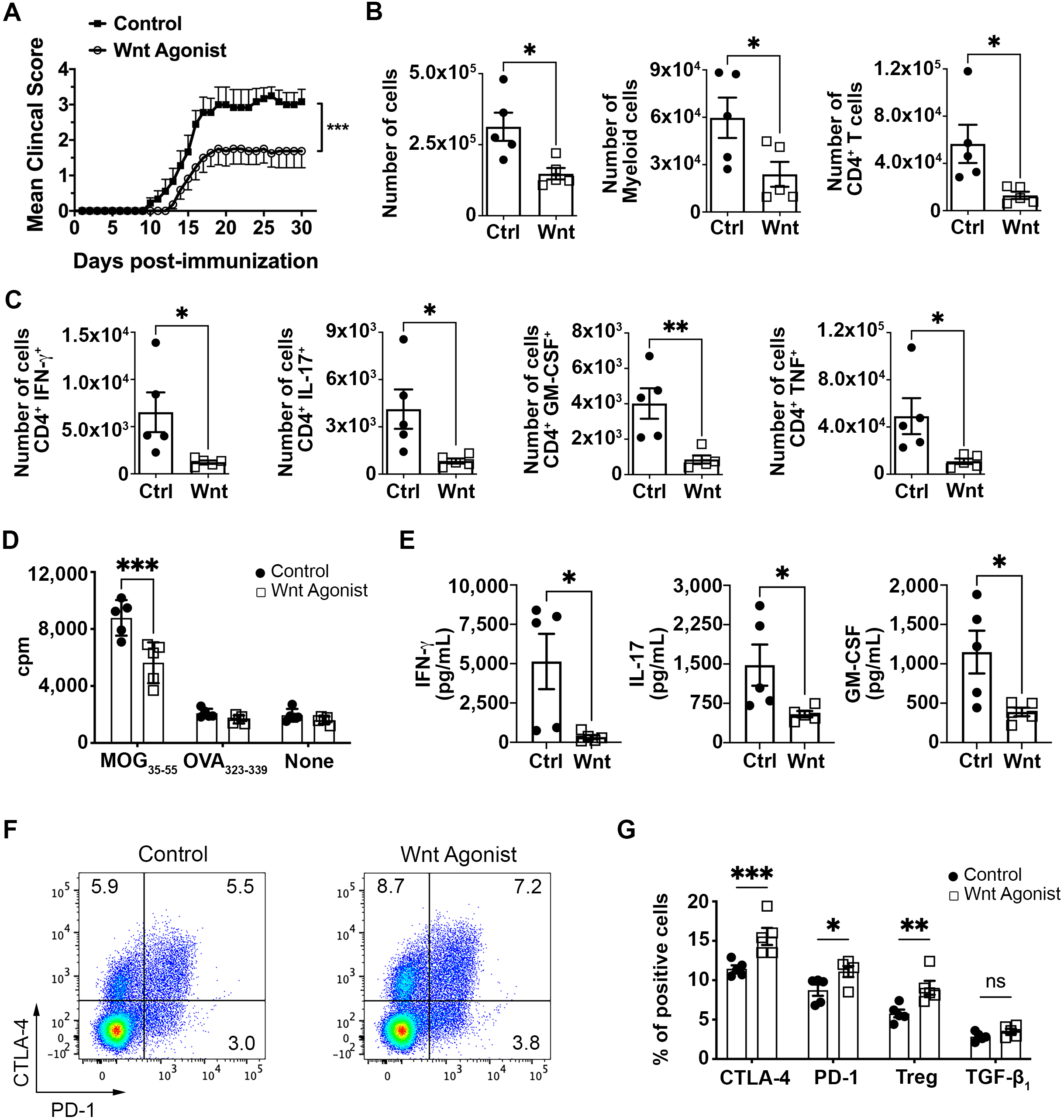
Prophylactic Wnt Agonist Treatment Ameliorates Chronic EAE. Active C-EAE was induced in C57BL/6 mice by immunization with MOG_35−55_. (**A**) Prophylactical treatment with Wnt agonist (2.5 mg/kg; open circles) significantly reduces EAE severity compared to the vehicle control (close squares) (*n* = 10 mice per group, \*\*\**p* < 0.001 by nonparametric Mann-Whitney test). (**B**) Flow cytometry quantification of total live cells, CD45^hi^ CD3^−^ CD11b^+^ infiltrating myeloid cells, CD45^hi^ CD11b^−^ CD3^+^ CD4^+^ T cells, and (**C**) IL-17, IFN-γ, GM-CSF and TNF-producing CD4^+^ T cells in the CNS of EAE mice treated with a control vehicle (closed circle) vs. a Wnt agonist (open square) at day 18 post-immunization (*n* = 5 mice per group; \**p* < 0.05, \*\**p* < 0.01 by Student’s *t* test). (**D**) Following prophylactic treatment, spleens were harvested at day 14 post-immunization and recall responses of total splenocytes (10^6^ cells/well) to MOG_35-55_ (20 μg/ml), OVA_323-339_ (20 μg/ml) or no peptide were measured after 72 hours of culture. Proliferation, determined via [^3^H]-TdR incorporation, and (**E**) cytokine production, measured by multiplex assay in response to MOG_35-55_, were reduced in the spleens of Wnt-treated mice (open square) compared to control-treated mice (closed circles) (*n* = 5 mice per group; \**p* < 0.05, \*\*\**p* < 0.001 by two-way ANOVA with a Bonferroni post-test and by Student’s *t* test). (**F**) Representative flow cytometry plots showing the expression of inhibitory molecules CTLA-4 and PD-1 on live CD4^+^ T cells following recall assays with MOG_35-55_. Data shown are representative of 5 independent experiments. (**G**) Compilation of CTLA-4, PD-1, TGF-β and Treg (CD25^+^ FoxP3^+^) expression by flow cytometry on live CD4^+^ T cells following recall assays (*n* = 5 mice per group; \**p* < 0.05, \*\**p* < 0.01, \*\*\**p* < 0.001 and ns = not significant by by two-way ANOVA with a Bonferroni post-test).

### Wnt agonist treatment reduces the inflammatory potential of myelin-reactive CD4^+^ T lymphocytes

To determine whether Wnt agonist treatment affected the antigen-specific T cell response, we isolated splenocytes from C-EAE mice on day 14 post-immunization and stimulated them with MOG_35-55_ peptide for 48 hours. Splenocytes from Wnt-treated mice exhibited significantly reduced recall responses, as determined by decreased proliferation and lower secretion of IFN-γ, IL-17 and GM-CSF (**Figure 5D and 5E**; n = 5 mice per group, \**p* < 0.05, \*\*\**p* < 0.001). These results confirm that Wnt agonist treatment inhibits effector cytokine production by T cells, supporting our *in vitro* data.

Our *in vitro* studies also revealed an upregulation of immunoregulatory molecules on T cells when co-culture with Wnt-activated APCs (**Figure 4C and Figure S6**). To assess whether this phenotype was reproduced *ex vivo*, we analyzed the T cells profile following the recall response. We found that following the recall assay, CD4^+^ T cells from Wnt-treated mice also presented a significant increase in CTLA-4 and PD-1 expression (**Figure 5F and 5G**, n = 5 mice per group, *\*p* < 0.05, \*\*\**p* < 0.001).

Unexpectedly, we also observed a higher percentage of Tregs following the recall response (**Figure 5G**, n = 5 mice per group, *\**\**p* < 0.01). Notably, this effect was previously detected only in our human *in vitro* studies, but not in our murine *in vitro* studies. No statistical difference was observed in TGF-β1 expression (**Figure 5G**).

### In vivo Wnt treatment promotes immune regulatory responses in myeloid cells

To confirm whether our *in vitro* data on myeloid cells held true *in vivo*, we analyzed the myeloid cell populations in the spleen, lymph nodes (LN) and CNS of treated mice, at day 12 post-immunization, one day after the final Wnt agonist injection.

We observed a significant upregulation of PD-L1 and PD-L2 expression by mDCs in all analyzed organs from Wnt agonist treated mice (**Figure 6A**; n = 6 mice per group, \**p* < 0.05, \*\**p* < 0.01). Additionally, macrophages in the spleen and CNS Wnt-treated mice exhibited increased expression of PD-L1 and PD-L2 (**Figure 6B**; n = 6 mice per group, \*\**p* < 0.01, \*\*\**p* < 0.001). Inflammatory monocytes also presented an upregulated expression of PD-L1 in the spleen and an increase expression of PD-L2 in the CNS of animals treated with the Wnt agonist (**Figure 6C**, n = 6 mice per group, \**p* < 0.05, \*\**p* < 0.01).

**Figure 6.**
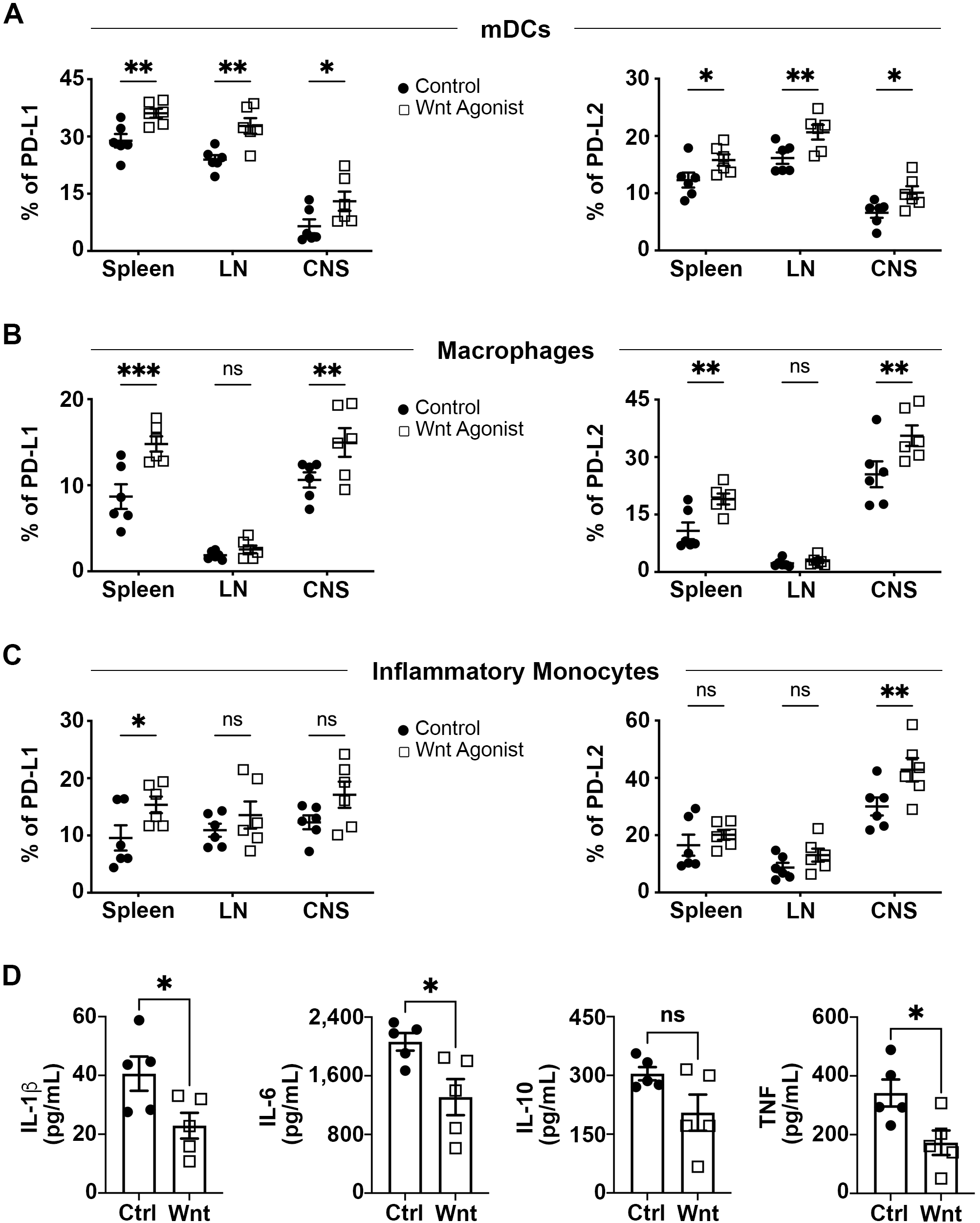
*In Vivo* Wnt Treatment Induces an Immunoregulatory Myeloid Cell Response. At day 14 post-immunization, spleens, lymph nodes (LNs) and homogenates of brain and spinal cords (CNS) from prophylactically treated C57BL/6 mice were harvested, and immune cells were isolated. Expression levels of inhibitory molecules PD-L1 and PD-L2 were assessed by flow cytometry on Wnt-treated EAE mice (open squares) and control EAE mice (closed circles) on (**A**) mDCs (Live^+^B220^−^CD11b^+^CD11c^+^Ly6G^−^), (**B**) macrophages (Live^+^B220^−^CD11b^+^CD11c^−^Ly6G^−^) and (**C**) inflammatory monocytes (Live^+^B220^−^CD11b^+^CD11c^−^Ly6C^hi^Ly6G^−^) (*n* = 5 mice per group; \**p* < 0.05, \*\**p* < 0.01, \*\*\**p* < 0.001 and ns = not significant by two-way ANOVA with a Bonferroni post-test). (**D**) Multiplex cytokine assay analysis of splenocytes 24 hours post LPS activation. Wnt-treated mice (open squares) show a significant decrease in IL-1β, IL-6 and TNF production, but no significant difference in IL-10 production compared to control EAE mice (closed circles) (*n* = 5, *p < 0.05 and ns = not significant by Student’s *t* test).

To further characterize the impact of Wnt signaling on myeloid cells, we analyzed their cytokine profile. Splenocytes from Wnt-treated mice showed a significant reduction in IL-1β, IL-6 and TNF production following LPS stimulation (**Figure 6D**; n = 5 mice per group, *\*p* < 0.05). However, no significant differences were observed in IL-10 secretion. Additionally, IL-23 and IL-12p70 levels were below the detection limit in most samples, preventing reliable analysis.

### Therapeutic Wnt treatment ameliorates disease in RR-EAE

We next investigated the therapeutic potential of the Wnt agonist in treating ongoing disease in PLP_139-151_-induced relapse remitting (RR)-EAE model in SJL/J mice. Mice were administered the Wnt agonist (2.5mg/kg) or vehicle control via i.p. injection 3 times per week, starting at the remission phase after the first relapse (day 34 post-immunization) and continued until the experiment’s conclusion (day 62 post-immunization).

Wnt agonist treatment significantly reduced relapse severity compared to vehicle-treated mice (**Figure 7A**; n = 8 mice per group, *\*\*\*p* < 0.001). This effect correlated with a reduction in CD45^hi^ CD3^−^ CD11b^+^ infiltrating myeloid cells, and CD45^hi^ CD3^+^ CD4^+^ T lymphocytes in the CNS of EAE mice sacrificed at day 62 post-immunization (**Figure S8**; n = 4 mice per group, *\*p* < 0.05, \*\**p* < 0.01).

**Figure 7.**
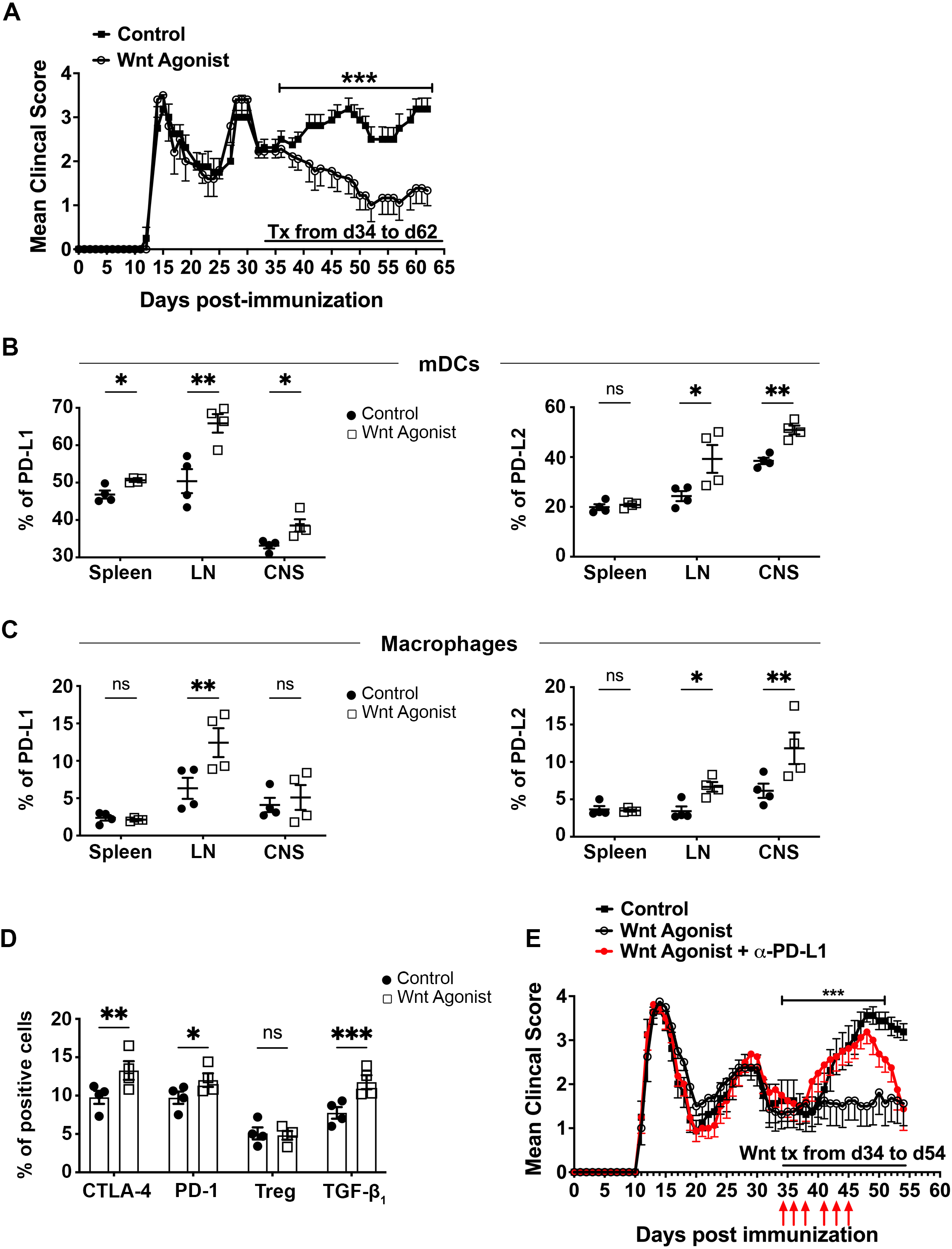
Therapeutic Wnt Treatment Reduces Relapse Severity in RR-EAE. EAE was induced in adult SJL mice by active immunization with PLP_139-151_. Wnt agonist (2.5 mg/kg) or vehicle control were injected i.p. 3 times a week starting at day 34 until day 62 post-induction. (**A**) Wnt agonist-treated mice (open circles) show a significant reduction in clinical scores compared to vehicle control-treated animals (closed squares) (*n*=8 mice per group; \*\*\**p* < 0.001 by nonparametric Mann-Whitney test). (**B**) At day 62 post-immunization, spleens, lymph nodes (LNs) and homogenates of brain and spinal cords (CNS) from therapeutically treated SJL/J mice were harvested, and immune cells were isolated. Expression levels of inhibitory molecules PD-L1 and PD-L2 were assessed by flow cytometry on Wnt-treated EAE mice (open squares) and control EAE mice (closed circles) on mDCs (Live^+^B220^−^CD11b^+^CD11c^+^Ly6G^−^), and (**C**) macrophages (Live^+^B220^−^CD11b^+^CD11c^−^Ly6G^−^) (*n* = 4 mice per group; \**p* < 0.05, \*\**p* < 0.01 and ns = not significant by two-way ANOVA with a Bonferroni post-test). (**D**) Following therapeutic treatment, spleens were harvested at day 62 post-immunization and recall responses of total splenocytes (10^6^ cells/well) to PLP_139-151_ (20 μg/ml) were assessed. Flow cytometry analysis of live CD4^+^ T cells was performed to evaluate the expression of CTLA-4, PD-1, TFG-β and Treg (CD25^+^ FoxP3^+^) (*n* = 4 mice per group; \**p* < 0.05, \*\**p* < 0.01, \*\*\**p* < 0.001 and ns = not significant by two-way ANOVA with a Bonferroni post-test). (**E**) EAE was induced in SJL mice by active immunization with PLP_139-151_. Wnt agonist (2.5 mg/kg; open squares) or the vehicle control (closed squares) were injected i.p. 3 times a week starting at day 34 until day 54 post-induction. A third group received Wnt treatment from day 34 to day 54 along with a PD-L1 blocking antibody (α-PD-L1; 10 mg/kg; closed red circles) from day 34 to day 45 (red arrows). (*n*=8 mice per group; \*\*\**p* < 0.001 by nonparametric Mann-Whitney test when comparing the Wnt agonist group to the Wnt agonist + α-PD-L1 from day 34 to day 45).

Moreover, Wnt agonist treated mice showed a significant reduction in CD4^+^ T cells expressing IL-17, IFN-γ, GM-CSF and TNF (**Figure S8**; n = 4 mice per group, *\*p* < 0.05, \*\**p* < 0.01). These findings provide the first evidence that Wnt agonist treatment can effectively control ongoing CNS inflammation and limit relapse severity in the RR-EAE model of MS, highlighting its potential therapeutic relevance.

To further explore the effects of Wnt agonist treatment in RR-EAE on myeloid cells in both the periphery and CNS, we analyzed their PD-L1 and PD-L2 expression. We found that mDCs from mice treated with the Wnt agonist showed an upregulated PD-L1 expression in the spleen, LN and CNS, and an upregulated expression of PD-L2 in the LN and CNS (**Figure 7B**; n = 4 mice per group, *\*p* < 0.05, \*\**p* < 0.01). Macrophages from Wnt agonist treated mice presented an upregulation of PD-L1 in the LN, and an upregulation of PD-L2 in the LN and CNS (**Figure 7C**; n = 4 mice per group, *\*p* < 0.05, \*\**p* < 0.01). However, inflammatory monocytes did not exhibit significant differences in PD-L1 or PD-L2 expression in any analyzed tissues (**Figure S9A**; n = 4 mice per group).

Cytokine analysis of myeloid cells revealed a significant reduction in IL-1β and TNF production in LPS-activated splenocytes from Wnt agonist treated mice (**Figure S9B**; n = 4 mice per group, *\*p* < 0.05, \*\**p* < 0.01). No significant differences in IL-6 and IL-10 secretion were observed. Once again, IL-23 and IL-12p70 were below detection limits, preventing a reliable analysis.

### Wnt agonist treatment reduces epitope spreading inflammatory potential of T cells

At day 62 post-immunization, we assessed encephalitogenic peptide-specific proliferation by stimulating splenocytes myelin peptides. Notably, splenocytes from Wnt agonist-treated mice exhibited decreased recall responses to both the disease-inducing PLP_139–151_ peptide and to PLP_178–191_, the dominant spread epitope, which would be responsible for the first relapse in RR-EAE (47) (**Figure S10A**; n = 4 mice per group, *p < 0.05, **p < 0.01). Furthermore, IFN-γ and GM-CSF production were significantly reduced in recall cultures with PLP_178-191_ (**Figure S10B**; n = 4 mice per group, *p < 0.05, **p < 0.01). These findings indicate that Wnt agonist treatment limits epitope spreading, aligning with its observed effects on disease progression.

Analysis of the T cell phenotype following recall culture with PLP_178-191_ further demonstrated an immunoregulatory shift, characterized by increased expression of CTLA-4, PD-1 and TGF-β1 in CD4^+^ T cells from Wnt agonist treated mice (**Figure 7D**; n = 4 mice per group, *\*p* < 0.05, \*\**p* < 0.01, *\*\*\*p* < 0.001).

### PD-L1 mediates the protective effect of Wnt agonist treatment

To determine whether the Wnt-induced regulatory myeloid cell phenotype was responsible for EAE relapse protection, we targeted PD-L1 since it is one of the most consistent upregulations observed in our human and mouse experiments. We designed an EAE experiment with 3 groups: 1) EAE mice treated with the Wnt agonist from day 34 to day 54; 2) EAE mice treated with the Wnt agonist from day 34 to day 54 and an anti-PD-L1 blocking antibody from day 34 to day 45; 3) EAE mice treated with vehicle control. As expected, Wnt agonist-treated mice were protected from new relapses (**Figure S11A**; n = 8 mice per group, *\*\*\*p* < 0.001). However, blocking PD-L1 abolished this protective effect, and these mice showed relapse severity comparable to control mice during the duration of anti-PD-L1 treatment (**Figure 7E**; n = 8 mice per group, *\*\*\*p* < 0.001). Interestingly, after discontinuing anti-PD-L1 injections at day 45, continued Wnt agonist treatment allowed mice to recover, and disease severity returned to levels observed in the Wnt agonist-treated group.

In contrast, when we repeated this experiment but blocked IL-10 instead of PD-L1, there was no effect on the protective role of Wnt agonist treatment (**Figure S11**; n = 8 mice per group). These findings demonstrate that the Wnt agonist exerts its protective effect primarily through PD-L1 upregulation, reinforcing its therapeutic potential in controlling CNS inflammation and relapse severity in RR-EAE.

## Discussion

In this study, we demonstrate that Wnt/β-catenin signaling reprograms myeloid cells into potent immunoregulators, suppressing inflammatory cytokines and reshaping CD4^+^ T cell responses. Our findings suggest that Wnt-activated monocytes and CD11b^+^ myeloid cells not only suppress inflammation but also actively promote immune tolerance, highlighting a crucial immunomodulatory role for Wnt signaling in both innate and adaptive immune responses.

This study provides the first direct *in vivo* evidence that Wnt activation in myeloid cells confers therapeutic benefits in an RR-EAE model. While previous studies have suggested an immunoregulatory role for Wnt in innate immune cells (48–50), our findings directly show that Wnt signaling modulates antigen presentation, suppresses autoreactive CD4^+^ T cell responses, and limits epitope spreading. This identifies a novel mechanism of immune regulation in neuroinflammation. Notably, while most MS therapeutics focus on direct inhibition of lymphocytes, our data suggest that targeting peripheral myeloid cells could be a powerful yet underutilized strategy.

Our results reinforce the growing evidence that Wnt/β-catenin signaling is a critical immune checkpoint in myeloid cells, driving tolerance in adaptive immunity and suppressing inflammation. Prior studies have shown that Wnt signaling skews dendritic cells toward a tolerogenic state, limiting their ability to stimulate effector T cell responses (34, 51). Similarly, Wnt/β-catenin activation in macrophages has been associated with reduced pro-inflammatory cytokine production and enhanced tissue repair (52, 53). Our study extends these findings by demonstrating that Wnt signaling in APCs actively suppresses CD4^+^ T cell inflammatory responses, further underscoring the pathway’s broader role in immune regulation.

The biological relevance of these findings is particularly significant in the context of autoimmune diseases such as MS. Given that excessive activation of pro-inflammatory T cells, particularly Th1 and Th17 subsets, drives MS pathology, our results suggest that targeting Wnt signaling in myeloid cells could restore immune balance. We showed that Wnt-activated APCs upregulate PD-L1 and PD-L2 expression, key regulators of lymphocyte activation and autoimmunity (54). Notably, PD-L1 expression on PBMCs is significantly reduced in MS patients compared to healthy controls (55), contributing to unchecked T cell activation and inflammation. Our findings suggest that Wnt activation restores myeloid cell immunoregulatory function by upregulating PD-L1, thereby promoting T cell tolerance. Increased PD-L1 expression was observed on APCs within both the CNS and secondary lymphoid organs, indicating that Wnt signaling enhances immune checkpoints at multiple levels. This effect was functionally significant, as PD-L1 blockade abrogated the protective effects of Wnt activation in RR-EAE, confirming that PD-L1 upregulation is a key mechanism by which Wnt signaling reduces inflammation. Further studies should investigate the specific contribution of PD-L2 in mediating the protective effects of Wnt signaling.

We also demonstrated that Wnt-activated myeloid cells downregulate proinflammatory cytokines essential for CD4^+^ T cell polarization into Th1 or Th17 subsets. IL-12 is required for differentiating naïve CD4^+^ T cells into the Th1 lineage (15), while IL-1β and IL-23 promote Th17 polarization (19). Notably, IL-23 is crucial for driving pathogenic Th17 cells (56). Our co-culture experiments confirmed that T cells cultured with Wnt-activated myeloid cells exhibited a significant reduction in cytokines associated with these subsets, further supporting the role of Wnt signaling in limiting pathogenic T cell responses.

Furthermore, Wnt-treated immune cells exhibited a diminished recall response to myelin peptide in both C- and RR-EAE models, indicating that Wnt signaling restricts epitope spreading. Given that epitope spreading contributes to disease progression in MS, our findings indicate that Wnt activation may not only modulate acute inflammation but also prevent the expansion of autoreactive T cell responses over time. Mechanistically, T cells from Wnt-treated mice showed upregulated expression of immune checkpoint molecules, including PD-1 and CTLA-4, both of which play key roles in inhibiting T cell activation (57). Additionally, given that PD-1 and CTLA-4 upregulation is also associated with T cell anergy (58), further investigation is needed to determine whether Wnt signaling directly induces an anergic state in autoreactive T cells. Furthermore, some of our other *in vivo* findings suggest that the canonical Wnt pathway may have direct effects on CD4^+^ T cells, a possibility we plan to explore in future studies.

Despite these promising insights, our study has some limitations. First, we used SB216763, a Wnt agonist that activates β-catenin signaling by inhibiting GSK-3β. SB216763 is widely used in research on the canonical Wnt pathway (59). However, it also has off-target effects on signaling pathways, including mTOR and NF-κB (60). Although we confirmed that SB216763 activates the canonical Wnt pathway, we cannot rule out potential contributions from other signaling cascades. To address this, future studies should compare the immunomodulatory effects of SB216763 with more selective Wnt activators, such as Wnt ligands, Frizzled receptors agonists, or direct β-catenin activators. Another key consideration is the potential risk associated with systemic Wnt activation. While our findings suggest that Wnt-induced immune modulation is beneficial in suppressing inflammatory responses, Wnt signaling has also been linked to tumorigenesis and fibrosis in certain contexts (61, 62). To mitigate these risks, future studies should explore cell-specific targeting strategies, such as nanoparticle-based delivery of Wnt activators to myeloid cells or engineered myeloid cells with controlled Wnt activation, or localized approaches like intrathecal administration to enhance therapeutic efficacy while minimizing unintended effects.

In summary, our study demonstrates that Wnt/β-catenin activation in myeloid cells promotes an immunoregulatory phenotype that suppresses pathogenic CD4^+^ T cell responses and limits neuroinflammation. By showing that Wnt signaling enhances immune tolerance through PD-L1 upregulation, reduces Th1/Th17 polarization, and restricts epitope spreading, our findings reveal a previously unrecognized mechanism for immune regulation in autoimmune disease. These results not only advance our understanding of myeloid cell biology in neuroinflammation but also highlights myeloid-targeted modulation as a promising therapeutic approach for MS and other inflammatory disorders. Given the current emphasis on lymphocyte-targeted therapies, our study underscores the potential of harnessing myeloid cell modulation as an alternative or complementary approach to restore immune homeostasis.

## Acknowledgments

We thank everyone in the Miller and Ifergan labs for their support. This work was supported by grants from the National Institutes of Health Grant R35GM146890 (II) and the National Multiple Sclerosis Society PP-2008-37129 (II).

## Author contributions

AA and II performed all the experiments of this study. SDM provided intellectual input throughout the project. The paper was written by AA, SDM and II.

