## Supplemental Information for "Wnt-Activated Immunoregulatory Myeloid Cells Prevent Relapse in Experimental Autoimmune Encephalomyelitis and Offer a Potential Therapeutic Strategy for Multiple Sclerosis"

**FIGURE LEGENDS**

**Figure S1. Wnt Activation Confirms β-Catenin Stabilization.** $CD14^+$ Monocytes were isolated from the blood of healthy donors and treated with either a Wnt agonist (SB216763, 10μM) or vehicle (DMSO) for 18 hours. A second stimulation with the Wnt agonist (10μM) or control vehicle (DMSO) was performed for 20min for ELISA quantification of total (left panel) and phosphorylated β-catenin (right panel) ($n$ = 6 donors, ***$p < 0.001$ and ns = not significant by paired Student's $t$ test).

**Figure S2. Activation of Wnt/β-Catenin Pathway Drives an Anti-Inflammatory Profile in Human Monocytes.** $CD14^+$ Monocytes were isolated from the blood of healthy donors and treated with either a Wnt agonist (SB216763, 10μM; right) or vehicle (DMSO; left) for 18 hours. Cells were then washed, and flow cytometry analysis was performed. Compilation of costimulatory (CD80, CD86, CD40), inhibitory (PD-L1, PD-L2) molecules and cytokines (IL-1b, IL-6, IL-10, IL-12p40, IL-23p19, TNF) expression ($n$ = 7 donors, *$p < 0.05$, **$p < 0.01$, ***$p < 0.001$ and ns = not significant by paired Student's $t$ test).

**Figure S3. Wnt/β-Catenin Activation Induces STAT Signaling in Human Monocytes.** $CD14^+$ Monocytes were isolated from the blood of healthy donors and treated with either a Wnt agonist (SB216763, 10μM) or vehicle (DMSO) for 18 hours. A second stimulation with the Wnt agonist (10μM) or control vehicle (DMSO) was performed for 20min for the ELISA quantification of phosphorylated STAT1 (pSTAT1), pSTAT3 (Y705), pSTAT3 (S727) and pSTAT6 (Y641) analysis ($n$ = 7 donors, **$p < 0.01$, ***$p < 0.001$ and ns = not significant by paired Student's $t$ test).

**Figure S4. Cytokine Secretion from Naïve and Memory CD4$^+$ T Cells in mixed leukocyte reaction.** Human CD4$^+$ T lymphocytes were co-cultured in a mixed leukocyte reaction (MLR) with Wnt-treated (SB216763, 10μM; open squares) or control (DMSO; closed circles) monocytes for 5 days. Multiplex cytokine assays were performed to measure secretion of IFN-γ, IL-17, GM-CSF, TNF, and IL-10 from the co-culture of monocytes with (**A**) memory CD4$^+$ CD45RA$^-$ T cell or with (**B**) naïve CD4$^+$ CD45RA$^+$ T cells ($n$ = 11 MLRs, *$p < 0.05$, **$p < 0.01$ and ns = not significant by paired Student's $t$ test).

**Figure S5. Murine Myeloid Cell Characterization Following Wnt Activation.** CD11b$^+$ splenocytes were isolated from healthy adult C57BL/6 mice and treated with either a Wnt agonist (SB216763, 10μM) or vehicle (DMSO) for 24 hours. Compilation of costimulatory (CD80, CD86), inhibitory (PD-L1, PD-L2) molecules and cytokines (IL-1b, IL-6, IL-10, IL-12p40, IL-23p19, TNF) expression ($n$ = 5 experiments, *$p < 0.05$, **$p < 0.01$ and ns = not significant by paired Student's $t$ test).

**Figure S6. Naïve CD4$^+$ T cells Profile After Co-Culture with Wnt-Activated CD11b$^+$ Myeloid Cells from 2D2 Mice.** Naïve CD4$^+$ CD62L$^{hi}$ CD44$^{lo}$ T cells isolated from the spleen of healthy adult 2D2 mice were co-cultured with either Wnt-activated (right) or vehicle (left) murine CD11b$^+$ cells for 4 days. Flow cytometry analysis of naïve CD4$^+$ T cell proliferation was assessed by % of Ki-67$^+$. CTLA-4, PD-1, Tregs (CD25$^+$ FoxP3$^+$), IL-17, IFN-γ, GM-CSF, TNF, TGF-β, and IL-10 expression were also assessed via flow cytometry ($n$ = 5 experiments, **$p < 0.01$ and ns = not significant by paired Student's $t$ test).

**Figure S7. Dose Response of Wnt Agonist Treatment in Chronic EAE.** Active C-EAE was induced in C57BL/6 mice by immunization with $MOG_{35-55}$ and treated prophylactically with a Wnt agonist at 0.5 mg/kg (green circles), 2.5 mg/kg (red triangles), and 10 mg/kg (blue diamonds). Only the 2.5mg/kg and the 10mg/kg doses significantly reduced EAE severity compared to the vehicle control (DMSO; black squares) ($n$ = 5 mice per group, *$p$ < 0.05 by nonparametric Mann-Whitney test).

**Figure S8. Therapeutic Wnt Treatment Reduces Immune Cell Infiltration and Inflammation in RR-EAE.** EAE was induced in adult SJL mice by active immunization with $PLP_{139\text{-}151}$. Wnt agonist (2.5 mg/kg) or vehicle control were injected i.p. 3 times a week starting at day 34 until day 62 post-induction. Flow cytometry quantification of total live cells, $CD45^{hi}$ $CD3^{-}$ $CD11b^{+}$ infiltrating myeloid cells, $CD45^{hi}$ $CD11b^{-}$ $CD3^{+}$ $CD4^{+}$ T cells, and IL-17, IFN-γ, GM-CSF and TNF-producing $CD4^{+}$ T cells in the CNS of EAE mice treated with a control vehicle (closed circle) vs. a Wnt agonist (open square) at day 62 post-immunization ($n$ = 4 mice per group; *$p$ < 0.05, **$p$ < 0.01 by Student's $t$ test).

**Figure S9. Inflammatory Monocytes Profile and Cytokine Production by Myeloid Cells Following Therapeutic Wnt Treatment in RR-EAE.** EAE was induced in adult SJL mice by active immunization with $PLP_{139\text{-}151}$. Wnt agonist (2.5 mg/kg) or vehicle control were injected i.p. 3 times a week starting at day 34 until day 62 post-induction. (**A**) At day 62 post-immunization, spleens, lymph nodes (LNs) and homogenates of brain and spinal cords (CNS) from therapeutically treated SJL/J mice were harvested, and immune cells were isolated. Expression levels of inhibitory molecules PD-L1 and PD-L2 were assessed by flow cytometry on Wnt-treated EAE mice (open squares) and control EAE mice (closed circles) on inflammatory monocytes ($Live^{+}B220^{-}CD11b^{+}CD11c^{-}Ly6C^{hi}Ly6G^{-}$) ($n$ = 4 mice per group; ns = not significant by two-way ANOVA with a Bonferroni post-test). (**B**) Multiplex cytokine assay analysis of splenocytes 24 hours post LPS activation. Wnt-treated mice (open squares) show a significant decrease in IL-1β and TNF production, but no significant difference in IL-6 and IL-10 production compared to control EAE mice (closed circles) ($n$ = 4, *p < 0.05, ***p* < 0.01 and ns = not significant by Student's *t* test).

**Figure S10. Recall Assays Following Therapeutic Wnt Treatment in RR-EAE.** Following therapeutic treatment, SJL/J spleens from Wnt-treated mice (open square) and spleen of control-treated mice (closed circles) were harvested at day 62 post-immunization and recall responses of total splenocytes ($10^6$ cells/well) to $PLP_{139-151}$ (20 μg/ml), $PLP_{178-191}$ (20 μg/ml), $MBP_{84-97}$ (20 μg/ml), $OVA_{323-339}$ (20 μg/ml) or no peptide were measured after 72 hours of culture. (**A**) Proliferation was determined via [$^3$H]-TdR incorporation, and (**B**) cytokine production was measured by multiplex assay in response to the various peptides ($n$ = 4, *$p$ < 0.05, **$p$ < 0.01 and ns = not significant by two-way ANOVA with a Bonferroni post-test).

**Figure S11. Beneficial Wnt Treatment in RR-EAE is Not Dependent on IL-10.** EAE was induced in adult SJL mice by active immunization with $PLP_{139-151}$. Wnt agonist (2.5 mg/kg) or vehicle control were injected i.p. 3 times a week starting at day 34 until day 54 post-induction. (**A**) Wnt agonist-treated mice (open circles) show a significant reduction in clinical scores compared to vehicle control-treated animals (closed squares) ($n$=8 mice per group; ***$p$ < 0.001 by nonparametric Mann-Whitney test). (**B**) A third group received Wnt treatment from day 34 to day 54 along with a IL-10 blocking antibody (α-IL-10; 10 mg/kg; closed blue circles) from day 34 to day 45 (blue arrows). ($n$=8 mice per group; ns = not significant by nonparametric Mann-Whitney test when comparing the Wnt agonist group to the Wnt agonist + α-IL-10 from day 34 to day 45).

**Table S1. List of genes, Fold Regulation and *p*-values of Wnt-treated Monocytes vs Control Monocytes.**

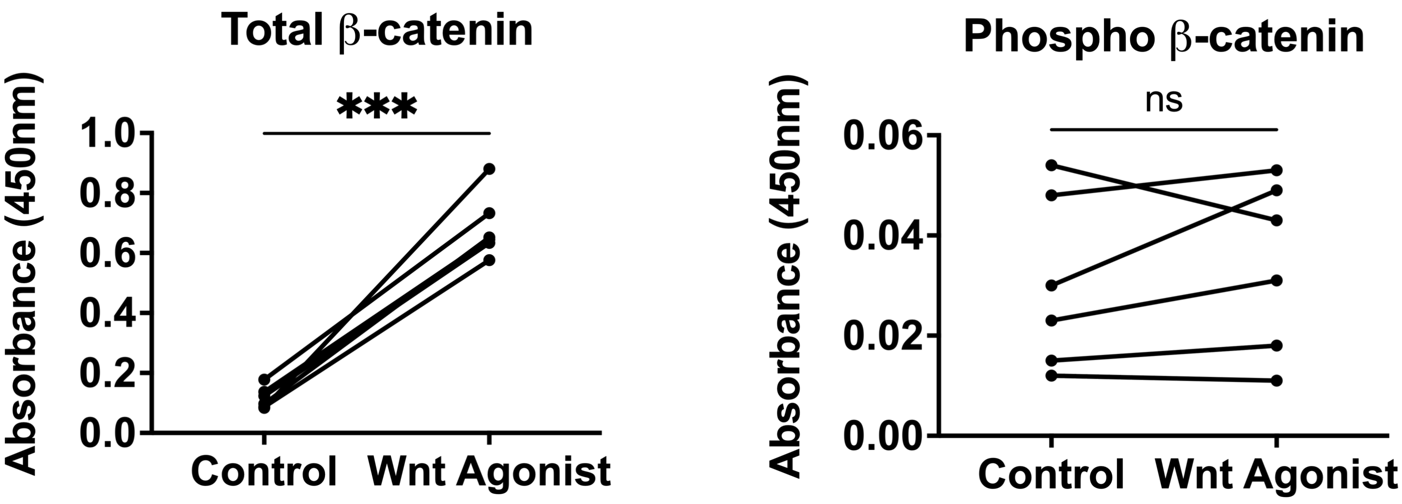

Total β-catenin
***
Absorbance (450nm)
1.0
0.8
0.6
0.4
0.2
0.0
Control
Wnt Agonist
Phospho β-catenin
ns
Absorbance (450nm)
0.06
0.04
0.02
0.00
Control
Wnt Agonist


Figure S1

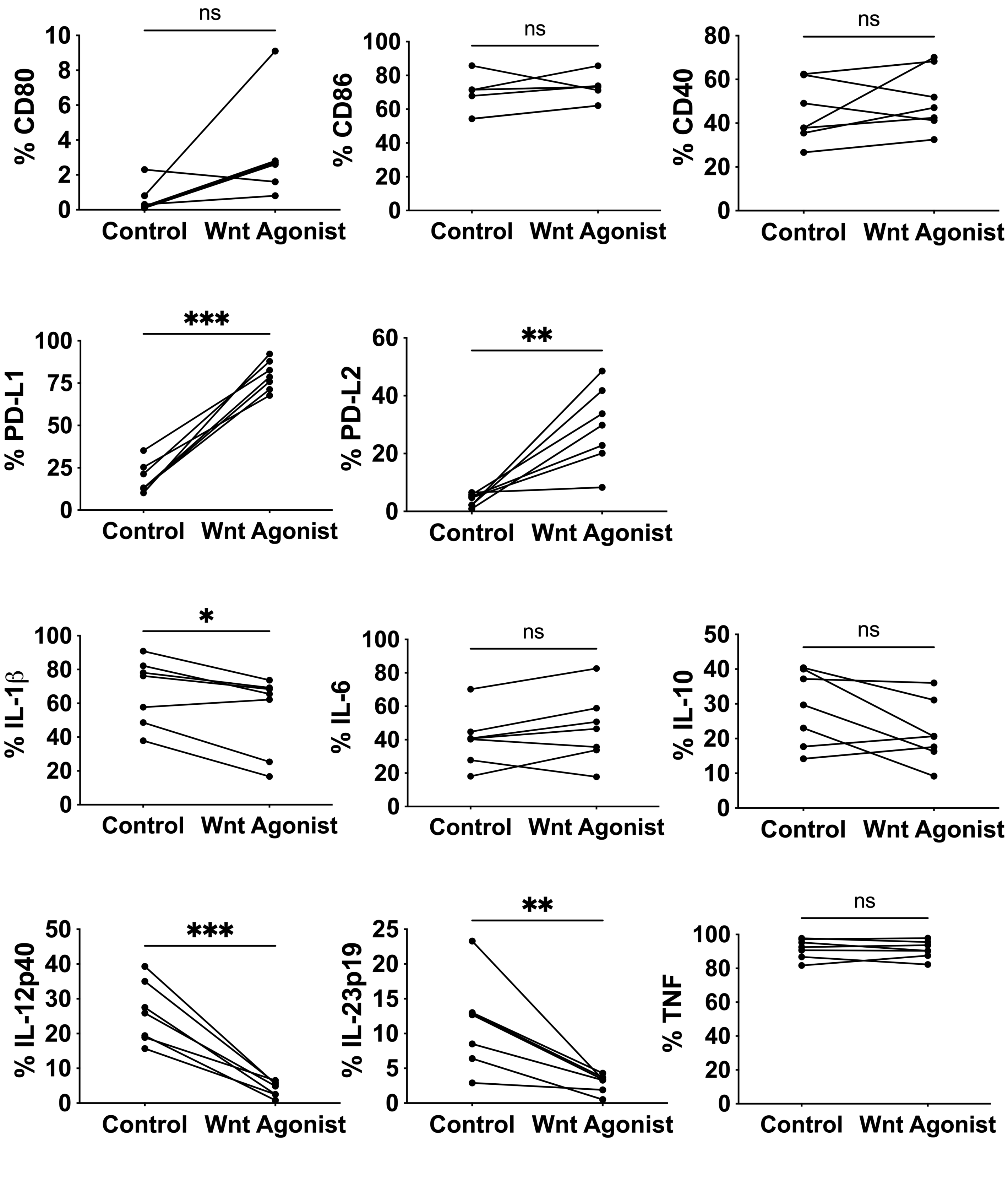
ns
% CD80
Control
Wnt Agonist
ns
% CD86
ns
% CD40
***
% PD-L1
**
% PD-L2
*
% IL-1β
ns
% IL-6
ns
% IL-10
***
% IL-12p40
**
% IL-23p19
ns
% TNF

Figure S2

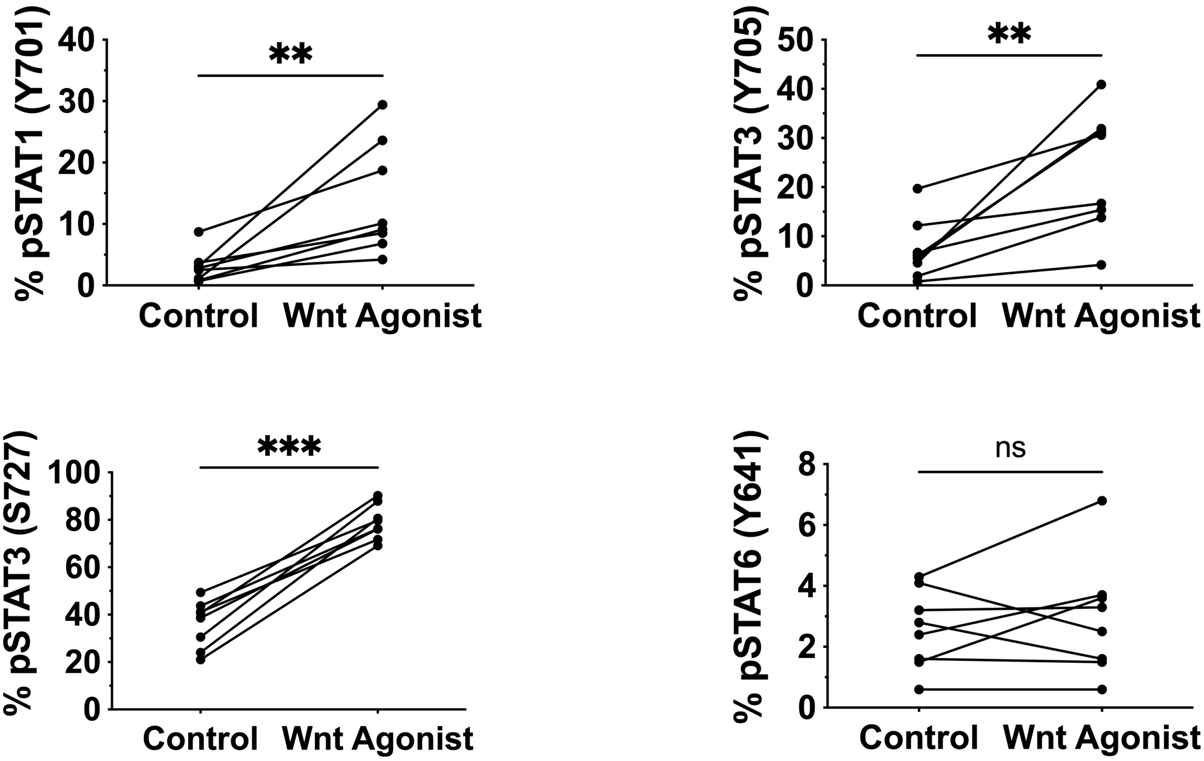
**
% pSTAT1 (Y701)
40
30
20
10
0
Control
Wnt Agonist
**
% pSTAT3 (Y705)
50
40
30
20
10
0
Control
Wnt Agonist
***
% pSTAT3 (S727)
100
80
60
40
20
0
Control
Wnt Agonist
ns
% pSTAT6 (Y641)
8
6
4
2
0
Control
Wnt Agonist

### Figure S3

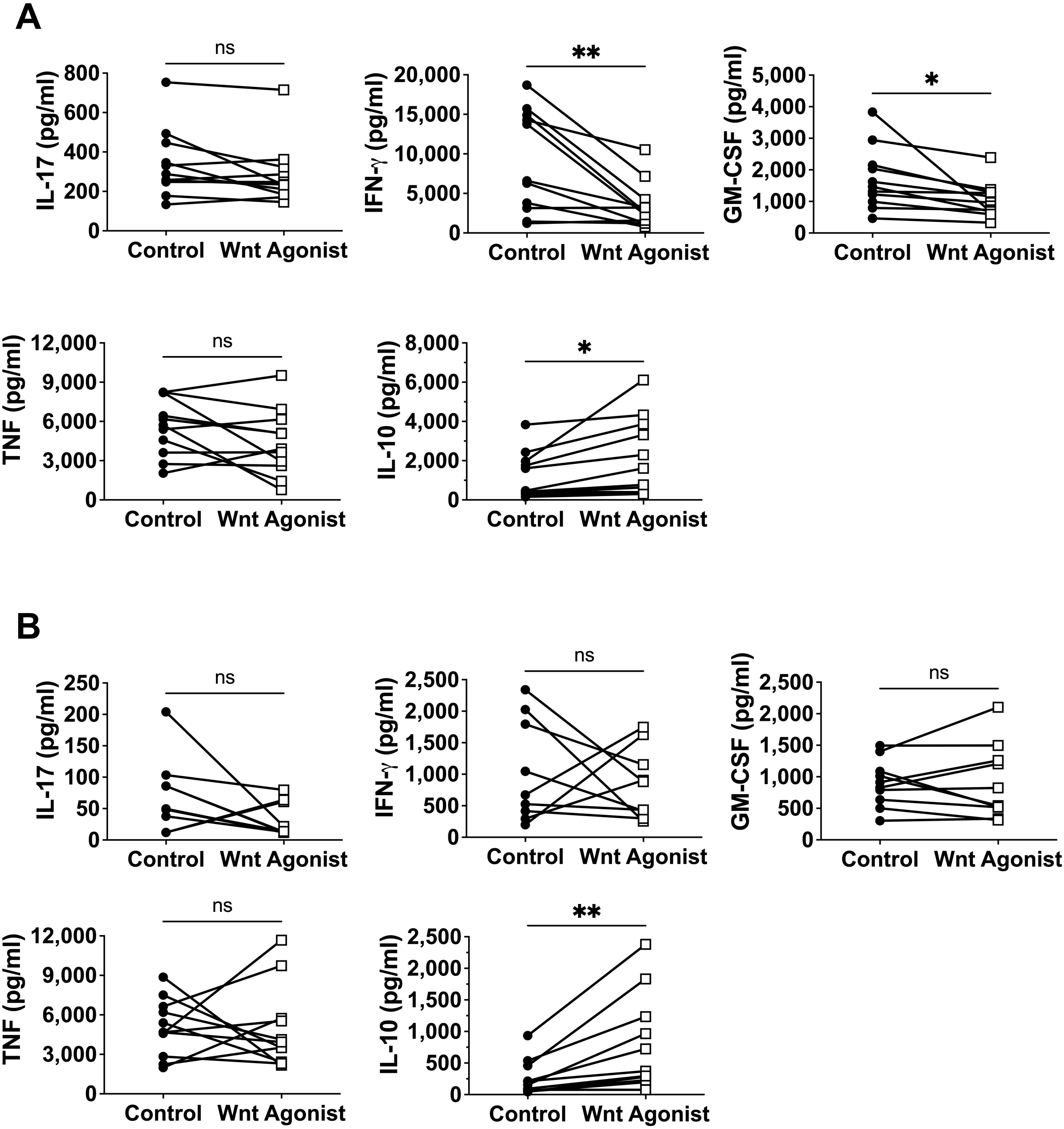
A
ns
IL-17 (pg/ml)
800
600
400
200
0
Control
Wnt Agonist
**
IFN-γ (pg/ml)
20,000
15,000
10,000
5,000
0
Control
Wnt Agonist
*
GM-CSF (pg/ml)
5,000
4,000
3,000
2,000
1,000
0
Control
Wnt Agonist
ns
TNF (pg/ml)
12,000
9,000
6,000
3,000
0
Control
Wnt Agonist
*
IL-10 (pg/ml)
8,000
6,000
4,000
2,000
0
Control
Wnt Agonist
B
ns
IL-17 (pg/ml)
250
200
150
100
50
0
Control
Wnt Agonist
ns
IFN-γ (pg/ml)
2,500
2,000
1,500
1,000
500
0
Control
Wnt Agonist
ns
GM-CSF (pg/ml)
2,500
2,000
1,500
1,000
500
0
Control
Wnt Agonist
ns
TNF (pg/ml)
12,000
9,000
6,000
3,000
0
Control
Wnt Agonist
**
IL-10 (pg/ml)
2,500
2,000
1,500
1,000
500
0
Control
Wnt Agonist

**Figure S4**

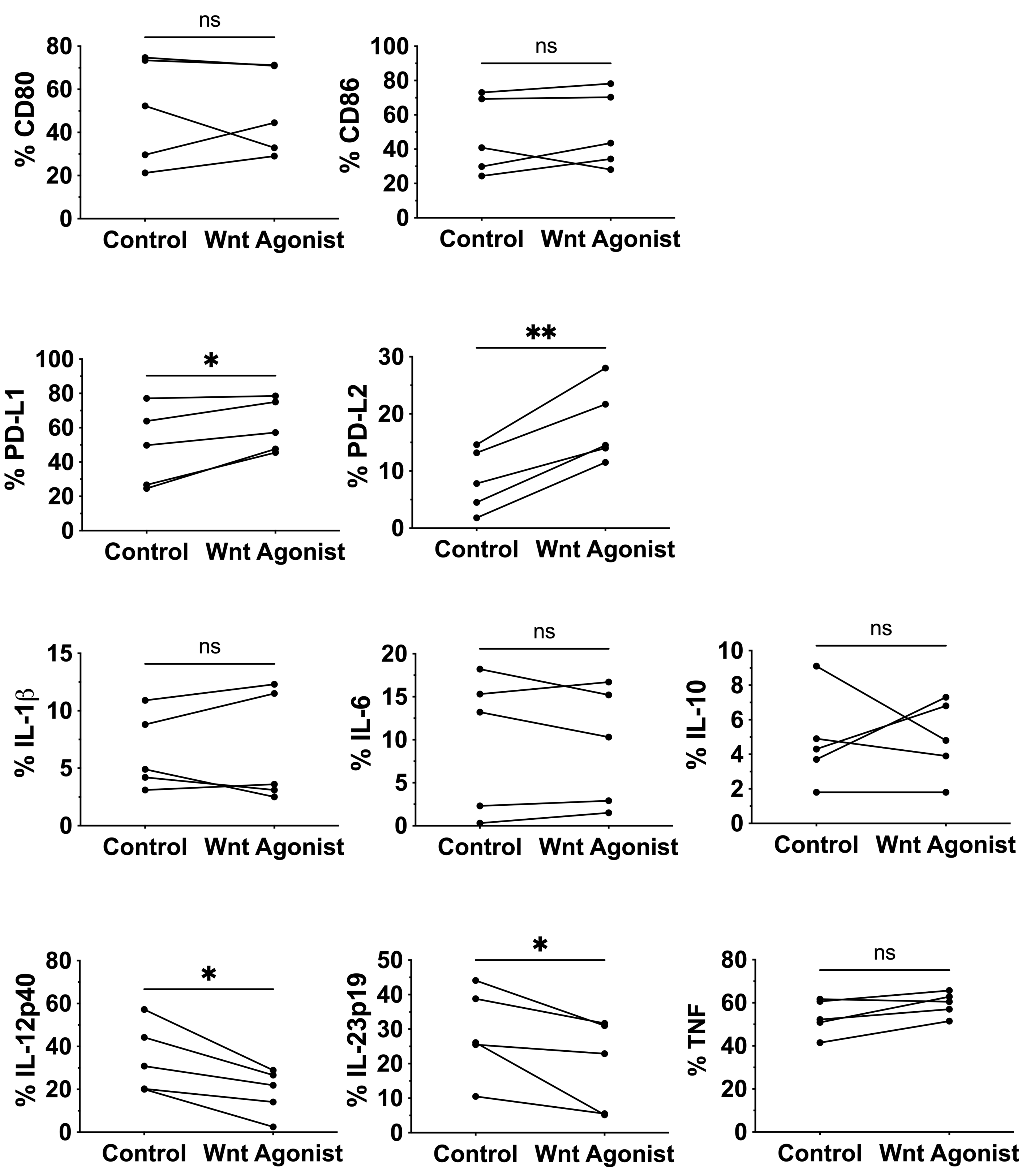
ns
% CD80
Control
Wnt Agonist
ns
% CD86
Control
Wnt Agonist
*
% PD-L1
Control
Wnt Agonist
**
% PD-L2
Control
Wnt Agonist
ns
% IL-1β
Control
Wnt Agonist
ns
% IL-6
Control
Wnt Agonist
ns
% IL-10
Control
Wnt Agonist
*
% IL-12p40
Control
Wnt Agonist
*
% IL-23p19
Control
Wnt Agonist
ns
% TNF
Control
Wnt Agonist

**Figure S5**

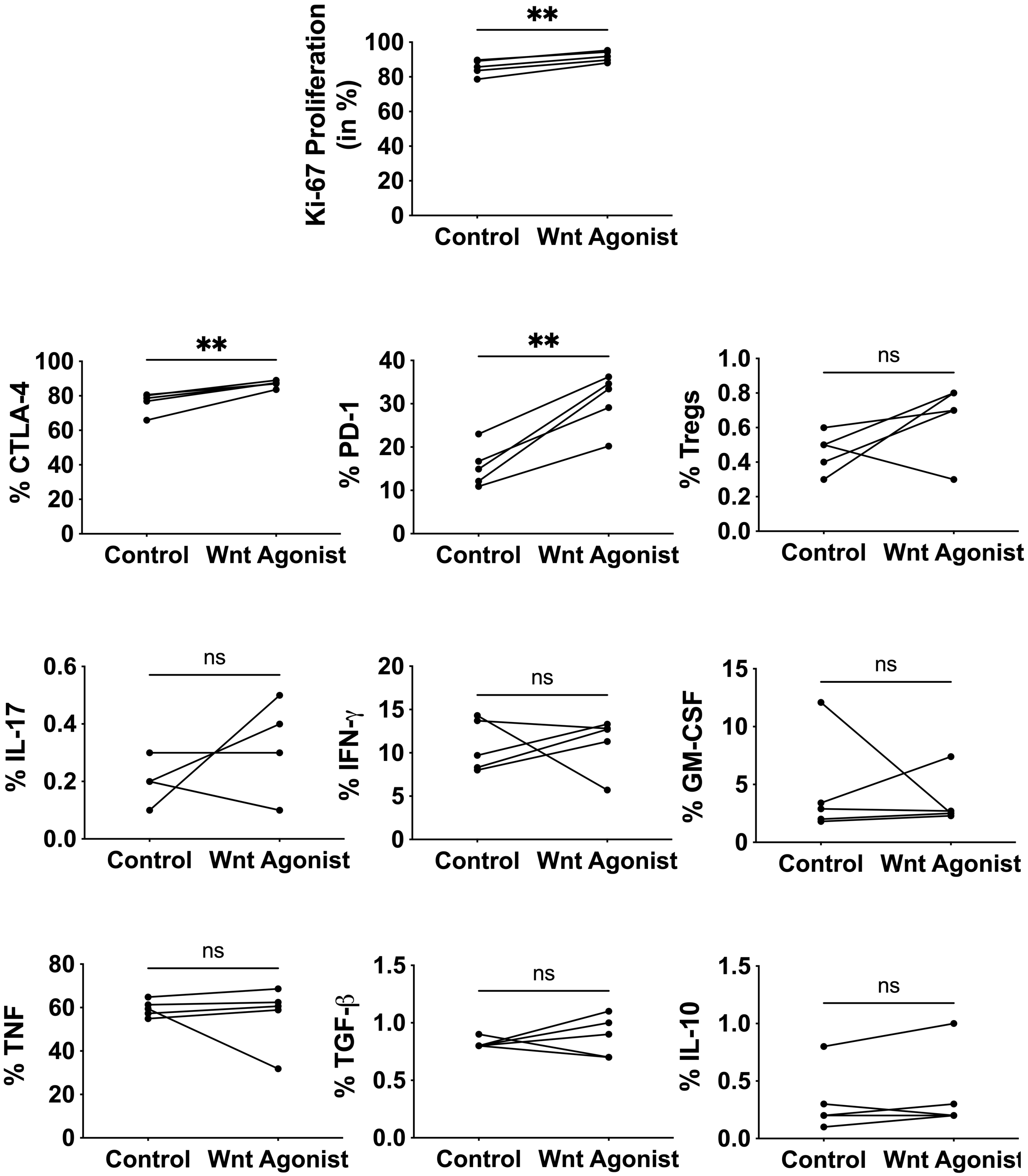
**
Ki-67 Proliferation (in %)
100
80
60
40
20
0
Control
Wnt Agonist
% CTLA-4
% PD-1
% Tregs
ns
% IL-17
% IFN-γ
% GM-CSF
% TNF
% TGF-β
% IL-10

### Figure S6

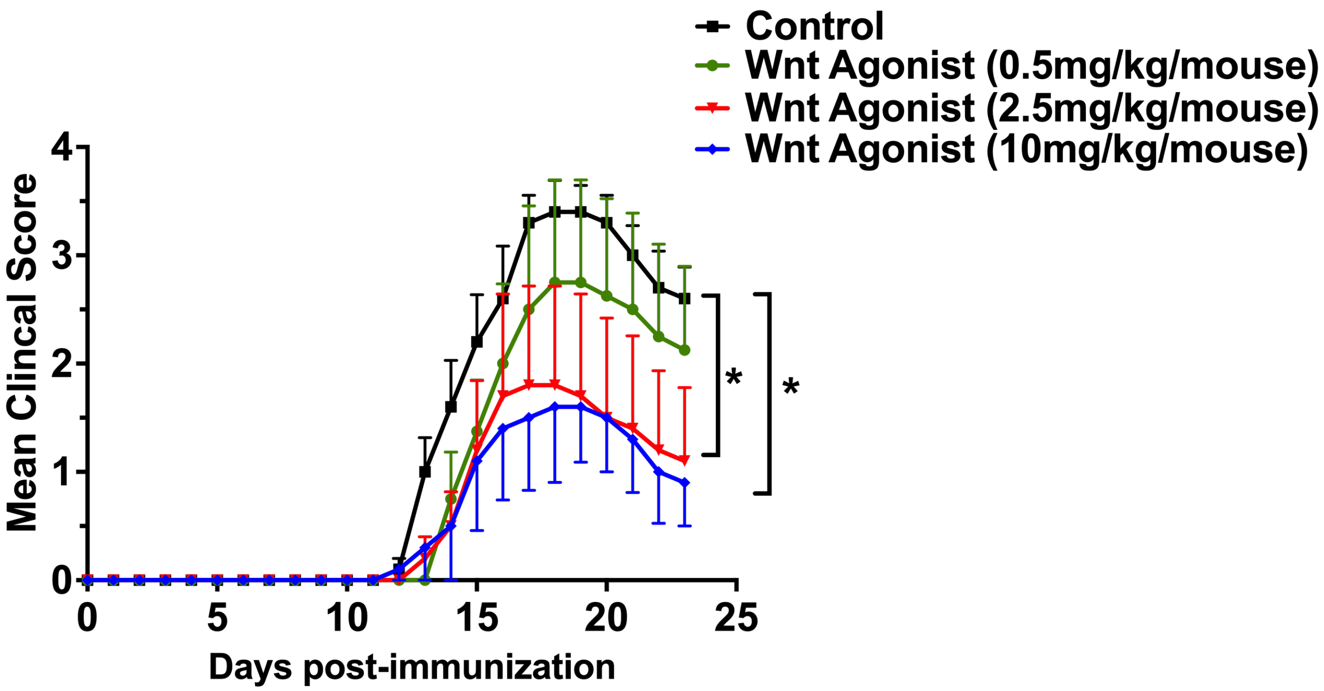
Control
Wnt Agonist (0.5mg/kg/mouse)
Wnt Agonist (2.5mg/kg/mouse)
Wnt Agonist (10mg/kg/mouse)
Mean Clincal Score
Days post-immunization
0
1
2
3
4
0
5
10
15
20
25
*
*

Figure S7

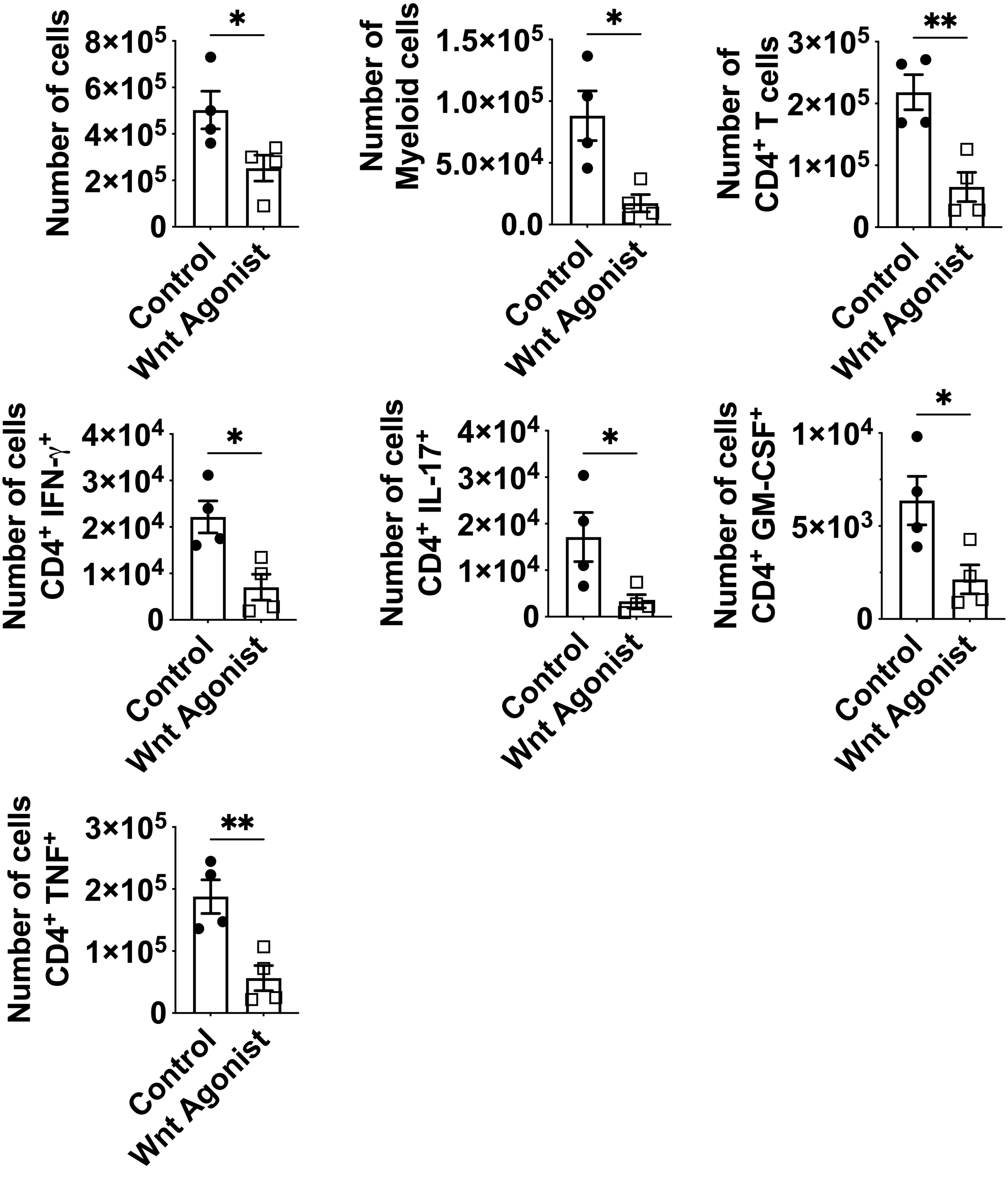
Number of cells
8×10⁵
6×10⁵
4×10⁵
2×10⁵
0
*
Control
Wnt Agonist
Number of Myeloid cells
1.5×10⁵
1.0×10⁵
5.0×10⁴
0.0
*
Control
Wnt Agonist
Number of CD4⁺ T cells
3×10⁵
2×10⁵
1×10⁵
0
**
Control
Wnt Agonist
Number of cells CD4⁺ IFN-γ⁺
4×10⁴
3×10⁴
2×10⁴
1×10⁴
0
*
Control
Wnt Agonist
Number of cells CD4⁺ IL-17⁺
4×10⁴
3×10⁴
2×10⁴
1×10⁴
0
*
Control
Wnt Agonist
Number of cells CD4⁺ GM-CSF⁺
1×10⁴
5×10³
0
*
Control
Wnt Agonist
Number of cells CD4⁺ TNF⁺
3×10⁵
2×10⁵
1×10⁵
0
**
Control
Wnt Agonist

Figure S8

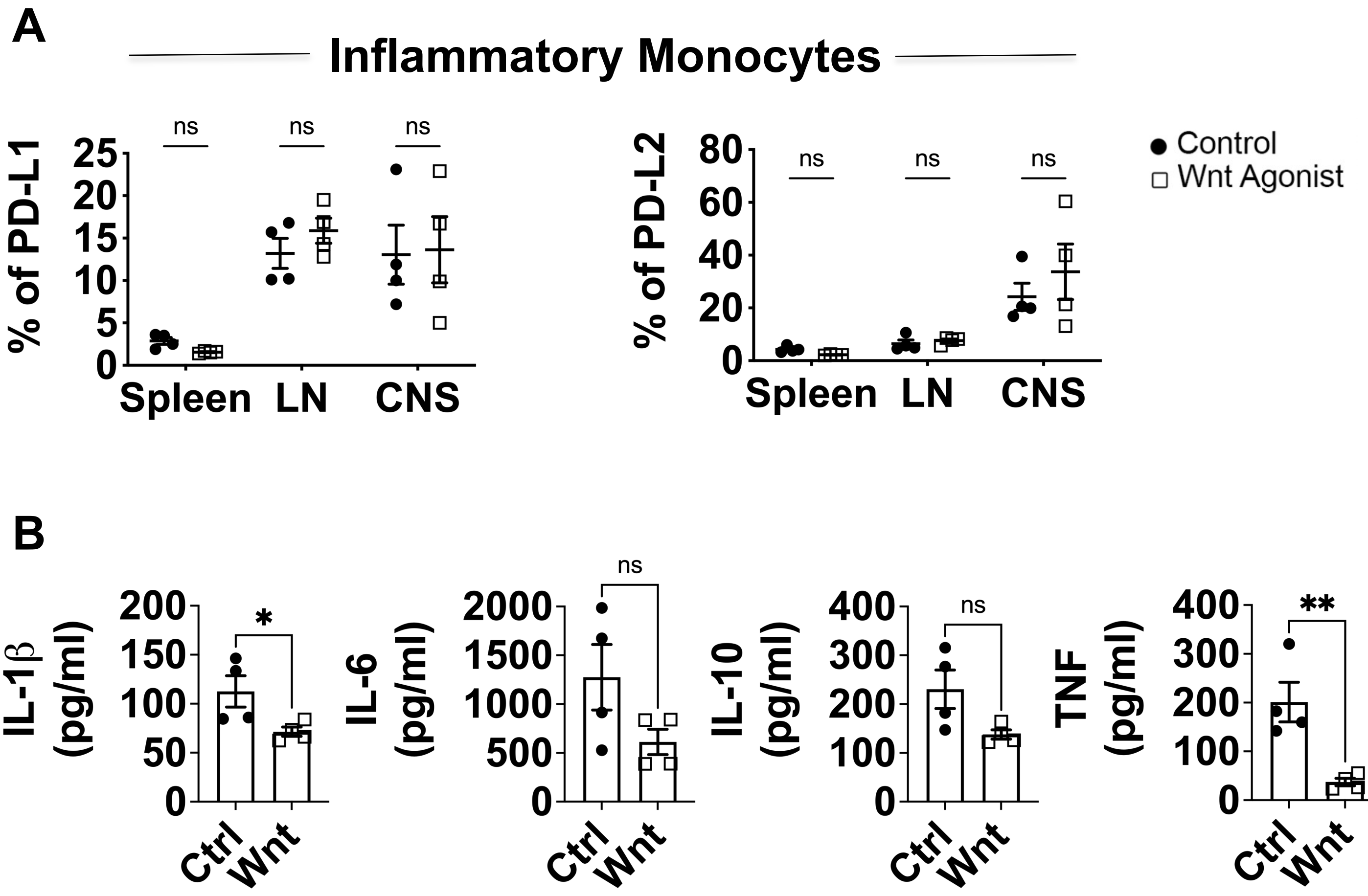
A
Inflammatory Monocytes
% of PD-L1
25
20
15
10
5
0
Spleen
LN
CNS
ns
% of PD-L2
80
60
40
20
0
Control
Wnt Agonist
B
IL-1β (pg/ml)
200
150
100
50
*
IL-6 (pg/ml)
2000
1500
1000
500
IL-10 (pg/ml)
400
300
TNF (pg/ml)
**
Ctrl
Wnt

Figure S9

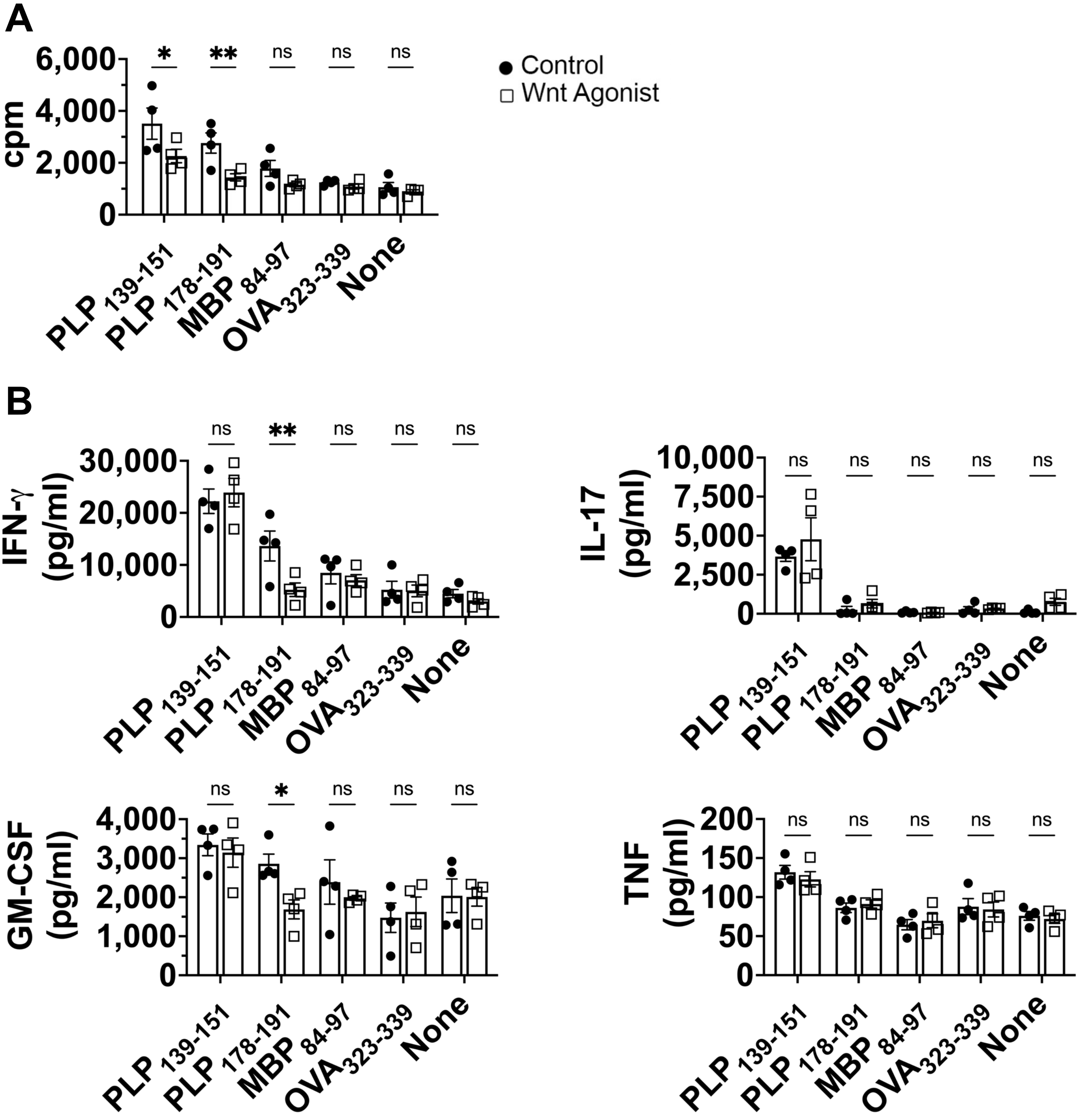
A
B
Control
Wnt Agonist
cpm
IFN-γ (pg/ml)
IL-17 (pg/ml)
GM-CSF (pg/ml)
TNF (pg/ml)
PLP 139-151
PLP 178-191
MBP 84-97
OVA323-339
None
ns
*
**

Figure S10

A

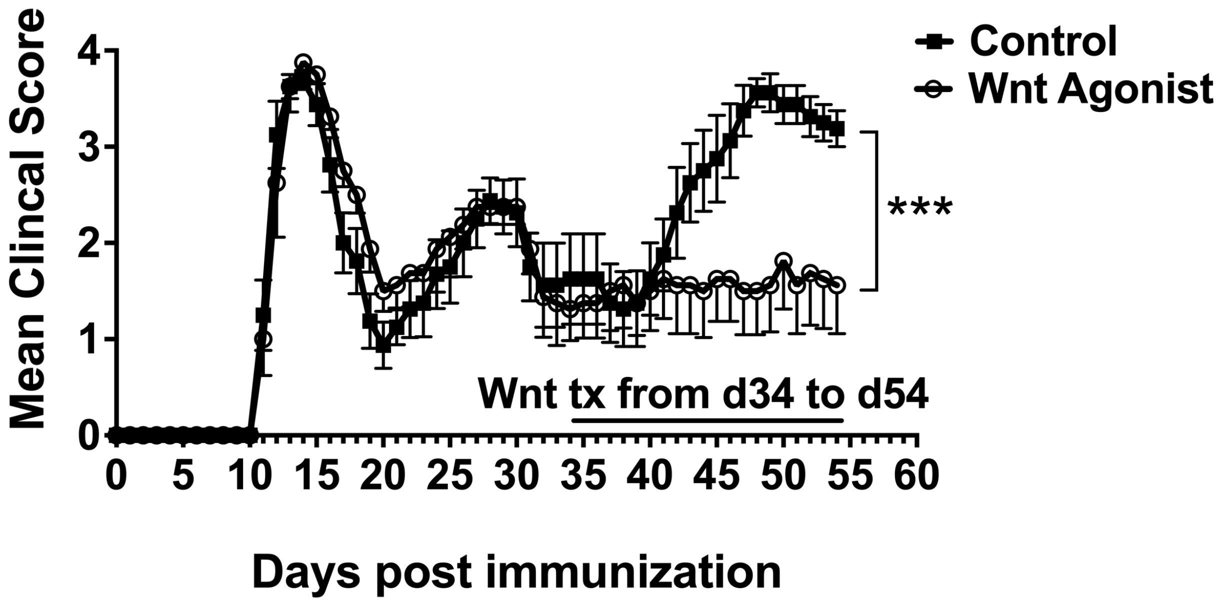


B

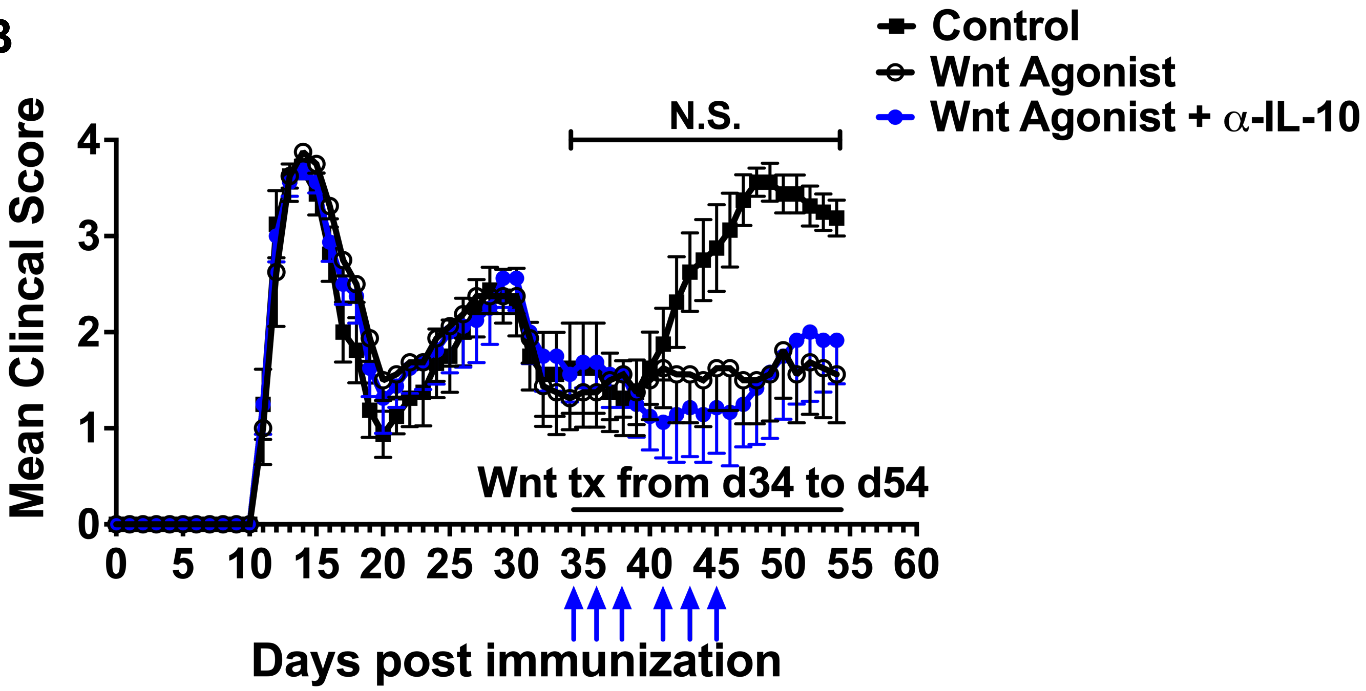


Figure S11

**Table S1: Genes, Fold Regulation and p-values of Wnt-treated Monocytes vs Control Monocytes**

| Gene Symbol | Fold Regulation | p-value |
|---|---|---|
| IL10RA | 6.53 | 0.0008085 |
| BCL2L1 | 5.03 | 0.00146998 |
| STAT1 | 30.23 | 0.00170649 |
| JAK2 | 8.38 | 0.00211843 |
| PRLR | 45.25 | 0.0027756 |
| JUN | 2.47 | 0.00370456 |
| SOCS2 | 11.22 | 0.00404039 |
| STAT5A | 2.33 | 0.00534233 |
| STAT2 | 12 | 0.00683697 |
| IRF1 | 147.28 | 0.00687429 |
| IL2RA | 600.81 | 0.01301517 |
| STUB1 | 2.15 | 0.01692617 |
| SRC | 10.02 | 0.01698045 |
| SPI1 | 3.73 | 0.01699484 |
| NOS2 | -7.73 | 0.0181667 |
| PTPRC | 4.54 | 0.02178822 |
| GRB2 | 2.35 | 0.02303477 |
| PTPN11 | 2.57 | 0.02698932 |
| OAS1 | 530.5 | 0.03012428 |
| JAK3 | 4.81 | 0.03079083 |
| STAT3 | 70.63 | 0.03132779 |
| FAS | 39.21 | 0.03653328 |
| PTPN1 | 2.49 | 0.03856096 |
| MCL1 | 17.77 | 0.04392335 |
| IL6ST | 3.31 | 0.04431181 |
| JAK1 | 15.97 | 0.04476332 |
| STAT4 | 7.61 | 0.04866687 |
| ISG15 | 120.57 | 0.04876576 |
| PIAS2 | 2.39 | 0.07216826 |
| TYK2 | 2.56 | 0.08911412 |
| SMAD2 | 39.13 | 0.10405927 |
| FCER2 | 16.26 | 0.1324951 |
| FCGR1A | 8.53 | 0.13345648 |
| SOCS3 | 2.74 | 0.16045172 |
| IL20 | 6.14 | 0.18611218 |
| F2 | 222.72 | 0.21486014 |
| NFKB1 | 3.5 | 0.22274323 |
| EPOR | 23.5 | 0.24225015 |

| | | |
|---|---|---|
| STAT6 | 347.96 | 0.2441007 |
| IL4R | 2.74 | 0.24459211 |
| SOCS4 | 2.06 | 0.26982519 |
| SMAD4 | 2.45 | 0.28737986 |
| JUNB | 22.73 | 0.30184692 |
| GATA3 | -5.05 | 0.31216504 |
| CRP | 1787.02 | 0.34637618 |
| USF1 | 2113.95 | 0.34647017 |
| EGFR | -59.53 | 0.34651015 |
| IL4 | -83.04 | 0.34659004 |
| PDGFRA | -8.02 | 0.34659365 |
| IFNG | -14.93 | 0.34659369 |
| SH2B1 | -11.17 | 0.3465943 |
| IRF9 | 2.83 | 0.34662132 |
| CRK | 2.36 | 0.34665607 |
| STAM | -4.38 | 0.34969064 |
| NR3C1 | 2.58 | 0.35073033 |
| IFNGR1 | 6.96 | 0.35366067 |
| CXCL9 | 4.2 | 0.36919286 |
| SMAD3 | 6.4 | 0.36921772 |
| INSR | 2.05 | 0.45825318 |
| CCND1 | 14.78 | 0.51325971 |
| CDKN1A | 7.39 | 0.52397972 |
| PIAS1 | -30.25 | 0.5710351 |
| OSM | -6.19 | 0.695129 |
| CEBPB | -10.41 | 0.76374167 |
| AKT1 | 76.67 | 0.77690087 |
| PIAS3 | 6.66 | 0.80403055 |
| LRG1 | 3.53 | 0.9140055 |
| SMAD1 | 2.28 | 0.94263693 |
| IFNAR1 | 138.47 | 0.94619735 |
